# Brown adipocyte NOSEMPE promotes nonmitochondrial thermogenesis and improves systemic metabolism through ATF4 activation

**DOI:** 10.1101/610220

**Authors:** Esther Paulo, Yun Zhang, Ruchi Masand, Tony L. Huynh, Youngho Seo, Danielle L. Swaney, Margaret Soucheray, David Jimenez-Morales, Nevan J. Krogan, Biao Wang

**Affiliations:** Cardiovascular Research Institute, Department of Physiology, University of California, San Francisco, San Francisco CA, 94158, USA; Department of Cellular and Molecular Pharmacology, University of California, San Francisco, San Francisco, CA 94158, USA; California Institute for Quantitative Biosciences, QBI, University of California, San Francisco, San Francisco, CA 94158, USA; J. David Gladstone Institutes, San Francisco, CA 94158, USA; Department of Radiology and Biomedical Imaging, University of California, San Francisco, CA 94143, USA; Department of Medicine, Division of Cardiovascular Medicine, Stanford University, Stanford, CA 94305, USA

**Author notes:** Corresponding Author: Biao Wang Ph.D., University of California, San Francisco.

## Abstract

Mitochondrial transcription factor A (Tfam)-mediated mtDNA maintenance and transcription, as well as leucine-rich PPR motif-containing protein (Lrpprc)-mediated mtRNA maturation and translation are essential steps of mtDNA-encoded electron transport chain (ETC) protein expression. ETC is essential for mitochondrial thermogenesis, the process of oxygen-dependent heat production inside the mitochondria in brown adipocytes. Here we describe that Tfam or Lrpprc deficiency in brown adipocytes cause <u>no</u>n-<u>s</u>ynchronized <u>E</u>TC <u>m</u>RNA and <u>p</u>rotein <u>e</u>xpression (NOSEMPE) and mitochondrial ETC imbalance, ultimately abolish mitochondrial thermogenesis. However, mice with NOSEMPE in brown adipocytes are cold resistant upon an acute 4°C cold challenge, because of augmented nonmitochondrial thermogenesis driven by the “NOSEMPE→ATF4→proteome turnover” pathway. Importantly, mice with either NOSEMPE or ATF4 overexpression in brown adipocytes are protected against high-fat-diet-induced metabolic abnormalities, indicating a positive association between nonmitochondrial thermogenesis in brown adipocytes and metabolic fitness. Thus, although brown adipocytes are defined by their unique ability to produce heat through mitochondrial respiration, our study demonstrates a novel cytosolic nonmitochondrial thermogenesis in brown adipocytes. Targeting this ATF4-dependent nonmitochondrial thermogenesis in brown adipocytes may represent a new therapeutic strategy for combating metabolic disorders.

---

Energy balance requires equivalent energy intake from food and energy expenditure for basal metabolism, physical activity and adaptive thermogenesis. The adaptive thermogenesis refers to the heat production in response to environmental changes, which mainly occurs in brown adipose tissue (BAT) containing specialized mitochondria-rich brown adipocytes ^1, 2^. Brown fat depots in humans have been recognized using 18F-fluoro-deoxyglucose positron emission tomography (^18^F-FDG PET) with computer-assisted tomography (CT), due to their higher glucose uptake activity. Particularly, brown fat ^18^F-FDG uptake activity gradually declines with aging and metabolic diseases ^3–6^. Thus, increasing brown fat abundance to boost adaptive thermogenesis has been proposed as a therapeutic strategy to offset the positive energy balance and to improve metabolic health in humans ^7, 8^.

The cold-induced adaptive thermogenesis (CIT), also called βAR-induced adaptive thermogenesis, is a primary source of heat production from brown adipocytes upon cold stimulation, which requires mitochondrial respiration and uncoupling protein 1 (Ucp1)-mediated uncoupling ^1, 9^. Numerous studies centered on cAMP- and peroxisome proliferator-activated receptor gamma coactivator 1-(PGC1-) dependent mitochondrial biogenesis have demonstrated that mitochondrial quantity control in brown adipocytes determines their thermogenic capacity ^9–11^.

Both nuclear DNA and mitochondrial DNA (mtDNA) encode subunits of electron transport chain (ETC); thus, the synchronization of nuclear- and mtDNA-encoded ETC protein expression is essential for mitochondrial respiration and then βAR-induced adaptive thermogenesis in brown adipocytes ^12–14^. We have recently described that brown adipocyte-specific mitochondrial transcription factor A (Tfam) knockout mice (Tfam^BKO^) exhibit a paradoxical trade-off between mitochondria-fueled βAR-induced adaptive thermogenesis and systemic metabolism in mice ^15^. In this study, we further demonstrate that disrupting mitochondrial quality (by the <u>no</u>n-<u>s</u>ynchronized <u>E</u>TC <u>m</u>RNA and <u>p</u>rotein <u>e</u>xpression, abbreviated as NOSEMPE) in brown adipocytes induces an ATF4-dependent nonmitochondrial thermogenesis that is instead fueled by proteome turnover in the cytosol. This nonmitochondrial thermogenesis in brown adipocytes can contribute to organismal thermoregulation under acute cold stress and promote metabolic health, which unveils a new function of brown adipocytes in systemic thermoregulation and metabolism.

## RESULTS

### Lrpprc regulates mtDNA ETC gene expression and mitochondrial respiration in brown adipocytes

Leucine-rich pentatricopeptiderepeat containing protein (LRPPRC) is a master regulator of mtDNA-encoded RNA maturation and stability (**Fig. 1a**) ^16–18^, and LRPPRC mutations cause cytochrome c oxidase (complex IV) deficiency in the French-Canadian variant of Leigh syndrome ^19–21^. We have generated the brown adipocyte-specific Lrpprc knockout (*Ucp1-Cre;Lrpprc^f/f^*, Lrpprc^BKO^) mice. Q-PCR and western blot confirmed that Lrpprc was efficiently deleted in the BAT of Lrpprc^BKO^ mice and not in other tissues (**Supplementary Fig.1a, b**). BAT thermogenic genes *Ucp1* and *Dio2* were reduced in these mice, although Ucp1 protein levels were not altered (**Supplementary Fig.1c, d**). Lrpprc deficiency induced “whitening” of brown adipocytes at room temperature (RT) (**Fig.1b**). But both wild-type and Lrpprc-deficient brown adipocytes exhibited unilocular morphology uniformly at thermoneutrality (30°C) (**Fig.1b**), where mitochondrial respiration in brown adipocytes is not needed for organismal thermoregulation.

**Figure 1.**
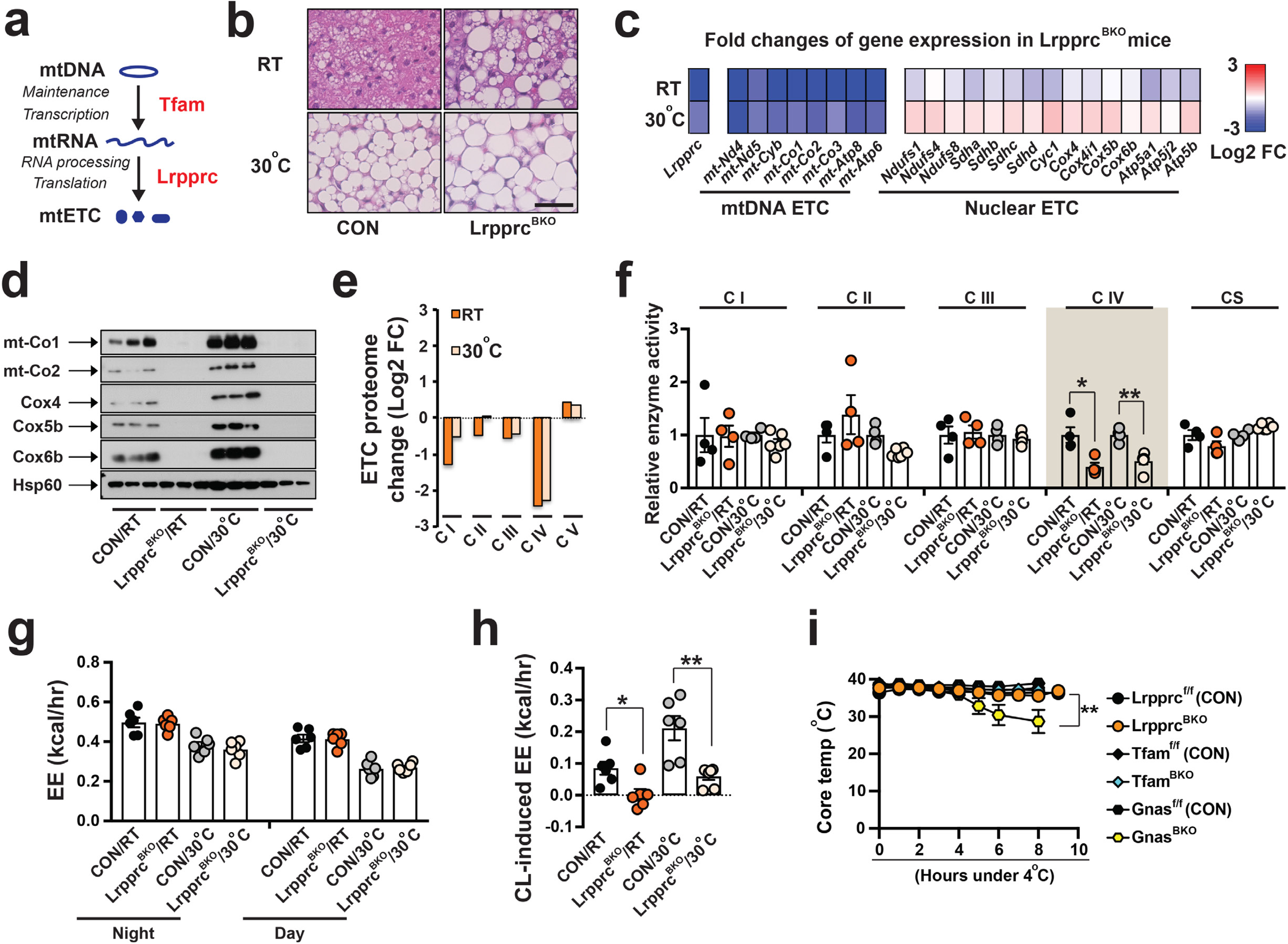
BAT NOSEMPE induces nonmitochondrial thermogenesis in Lrpprc^BKO^ mice. **(a)** Roles of Tfam and Lrpprc in mtDNA ETC protein expression. **(b)** Representative H&E staining of BAT from ∼8-12-week of male CON (*Lrpprc^f/f^*) and Lrpprc^BKO^ (*Ucp1-Cre;Lrpprc^f/f^*) mice housed at room temperature (RT) and thermoneutrality (30°C). Scale bar: 50μm. **(c)** Heatmap showing the log2 fold changes of *Lrpprc*, mtDNA-encoded and nuclear-encoded ETC subunits in the BAT from ∼8-12-week of male CON and Lrpprc^BKO^ mice housed at RT and 30°C. Sample size: CON/RT (n=4), Lrpprc^BKO^/RT (n=4), CON/30°C (n=4) and Lrpprc^BKO^/30°C (n=6). **(d)** Immunoblots of complex IV subunits (mt-Co1, mt-Co2, Cox4, Cox5b, Cox6b) and Hsp60 in isolated mitochondria from above mice. **(e)** Log2 fold change values of each proteome from BAT of Lrpprc^BKO^ mice. **(f)** Relative *in vitro* enzyme activities of Complex I to IV and citrate synthase (CS) in BAT of ∼8-12-week old male CON and Lrpprc^BKO^ mice housed at RT and 30°C. Sample size: CON/RT (n=4), Lrpprc^BKO^/RT (n=4), CON/30°C (n=4) and Lrpprc^BKO^/30°C (n=6). Average night and day EE **(g)** and hourly CL-induced EE **(h)** in ∼8-12-week old male CON and Lrpprc^BKO^ mice housed at RT and 30°C. Sample size: CON/RT (n=6), Lrpprc^BKO^/RT (n=6), CON/30°C (n=6) and Lrpprc^BKO^/30°C (n=7). **(i)** Cold tolerance test (CTT) of ∼8-12-week old male and female Lrpprc^BKO^, Tfam^BKO^, Gnas^BKO^ and their relative control mice housed at RT. Note: data from Gnas^BKO^ and relative control was from our previous publication ^24^. Sample size: Lrpprc^f/f^ (CON) (n=6), Lrpprc^BKO^ (n=9), Tfam^f/f^ (CON) (n=4), Tfam^BKO^ (n=4), Gnas^f/f^ (CON) (n=12), and Gnas^BKO^ (n=9). Data was presented as average ± SEM. Student t-test. *: p<0.05 and **: p<0.01.

As we expected, steady-state mRNA levels of most mtDNA-encoded genes were reduced in the BAT of Lrpprc^BKO^ mice, without significant changes in nuclear-encoded ETC genes or mtDNA copy numbers (**Fig.1c, Supplementary Fig.1e**). This specific reduction of mtDNA-encoded ETC genes was also observed in the BAT of Lrpprc^BKO^ mice housed at 30°C (**Fig.1c**). Immunoblots further confirmed that mtDNA-encoded complex IV proteins, mtCo1 and mtCo2, were reduced in isolated BAT mitochondria from Lrpprc^BKO^ mice (**Fig.1d, Supplementary Fig.2g**). Interestingly, nuclear-encoded complex IV subunits, such as Cox4, Cox5b and Cox6b, were also reduced at both ambient temperatures, even though their mRNA levels were largely unaffected (**Fig.1c, d**). In contrast, Atp5a (complex V), Uqcrc2 (complex III), Sdhb (complex II) and Ndufb8 (complex I) protein levels remain unaffected (**Supplementary Fig.2g**).

In order to obtain a global view of their mitochondrial proteome, we performed mass spectrometry analysis of freshly isolated BAT mitochondria from control and Lrpprc^BKO^ mice at normal chow housed at both RT and 30°C. We identified approximately 670 mitochondrial proteins for further analyses (**Supplementary Fig.2a-c, Supplementary Table 1**). The mass spectrometry analysis of mitochondrial proteome did not reveal profound metabolic reprogramming in Lrpprc-deficient brown adipocytes, including the enzymes involved in glycolysis, TCA cycle and beta-oxidation, except for the upregulation of Acot2 (**Supplementary Fig.2f**). Lrpprc deficiency did cause similar changes in mitochondrial ETC proteome at both RT and 30°C. Gene Ontology enrichment analysis showed that mitochondrial respiratory chain, NADH dehydrogenase (complex I) and cytochrome C oxidase (complex IV) activities were the most affected by Lrpprc deficiency at both ambient temperatures (**Supplementary Fig.2d, e**). The complex IV enzyme activity *in vitro* was attenuated in the BAT of Lrpprc^BKO^ mice (**Fig.1f**). To quantitate the BAT ETC proteome, we calculated the average log2 fold change (log2FC) values for all complex proteins identified and we found that complex IV was the most affected (**Fig.1e**), suggesting that an ETC proteome imbalance is induced by Lrpprc deficiency in brown adipocytes.

Previously we reported the BAT mitochondrial proteome in brown adipocyte-specific Lkb1 and Tfam knockout mice (Lkb1^BKO^ and Tfam^BKO^) ^15^. Clustering analysis showed that the mitochondrial proteomic changes induced by Tfam or Lrpprc deficiency were similar (**Supplementary Fig.2h**). Comparing the BAT mitochondrial proteome in Tfam^BKO^ and Lrpprc^BKO^ mice, we found that proteins involved in mitochondrial protein import (Timm10, Pam16, and Dnajc15) and proteases (Afg3l1, Afg3l2 and Lonp1) were selectively upregulated (**Supplementary Fig.2i**), suggesting that Lrpprc or Tfam deficiency induces a similar mitochondrial proteome remodeling in response to ETC proteome imbalance.

### NOSEMPE in brown adipocytes induces a nonmitochondrial thermogenesis

BAT adaptive thermogenesis is essential for organismal thermoregulation in rodents ^1, 2^. Genetic ablation of BAT in mice leads to increased cold sensitivity in an acute 4°C cold tolerance test (CTT) ^22, 23^, a rapid decrease of core body temperature (∼10°C decrease within several hours). Impaired mitochondrial biogenesis (in Gnas^BKO^ and betaless mice ^24, 25^) or Ucp1 deficiency (in Ucp1 knockout mice ^26^) in brown adipocytes also causes defective βAR-induced adaptive thermogenesis and increased cold sensitivity in CTT. Since mitochondrial respiration fuels βAR-induced adaptive thermogenesis in BAT ^1^, we then used indirect calorimetry experiments to measure βAR-induced adaptive thermogenesis, as well as basal energy expenditure (EE, calculated from oxygen consumption), respiratory exchange ratio (RER), food intake and physical activity of 8-10-week-old male control and Lrpprc^BKO^ mice at RT and 30°C. Body weights of Lrpprc^BKO^ and control mice were not different at this age. At basal state, there were no differences in basal EE, RER, food intake and physical activity (**Fig.1g, Supplementary Fig.3**). However, β3 agonist CL 316,423 (CL) stimulation significantly induced heat production in control mice, and this effect was absent in Lrpprc^BKO^ mice (**Fig.1h**). Consistently, BAT ^18^F-FDG uptake was also reduced in the Lrpprc^BKO^ mice (**Supplementary Fig.4a, b**).

Although both Tfam^BKO^ and Lrpprc^BKO^ mice did not have respiration-capable mitochondria in brown adipocytes and lacked βAR-induced adaptive thermogenesis ^15^, they were cold resistant during CTT paradoxically (**Fig.1i**). This was in sharp contrast to the Gnas^BKO^ mice, which also lacked βAR-induced adaptive thermogenesis but were cold sensitive during CTT ^24^. We reason that Lrpprc-deficiency, like Tfam-deficiency, causes a specific reduction of mtDNA-encoded ETC gene and protein expression (**Fig.1a**) ^20^, and consequently the <u>no</u>n-<u>s</u>ynchronized <u>E</u>TC <u>m</u>RNA and <u>p</u>rotein <u>e</u>xpression (NOSEMPE for short) ^12^. NOSEMPE specifically disrupts mitochondrial quality, not quantity. Although defective mitochondrial quantity and quality in BAT equally abolish βAR-induced adaptive thermogenesis, NOSEMPE might induce a compensatory thermogenic process that occurs outside the mitochondria, a hypothetical “nonmitochondrial thermogenesis”. In comparison, the conventionally viewed cold-induced, Ucp1-mediated heat production inside mitochondria in brown adipocytes is referred to mitochondrial thermogenesis hereafter.

### NOSEMPE activates ATF4-ISR in brown adipocytes

In order to identify the underlying mechanisms for NOSEMPE-induced nonmitochondrial thermogenesis, we first performed RNA-seq experiments from the BAT of male control and Lrpprc^BKO^ mice housed at RT and 30°C. Clustering analysis of differentially expressed genes (DEGs) suggested that thermoneutral housing affected BAT transcriptome more profoundly than Lrpprc deficiency (**Fig.2a**). Volcano plots of DEGs showed that there were approximately 6 times more up- and down-regulated DEGs in Lrpprc^BKO^ mice at 30°C (**Fig.2b**). Within downregulated DEGs, 132 genes were commonly observed at both RT and 30°C. The Oxidative phosphorylation pathway was the most significantly enriched in downregulated DEGs at both ambient temperatures (**Supplementary Fig.5a, b**). Indeed, the sequencing reads of all 13 mtDNA-encoded ETC subunits were reduced in the BAT of Lrpprc^BKO^ mice (**Supplementary Fig.5c**), consistent with our q-PCR analysis (**Fig.1c**).

**Figure 2.**
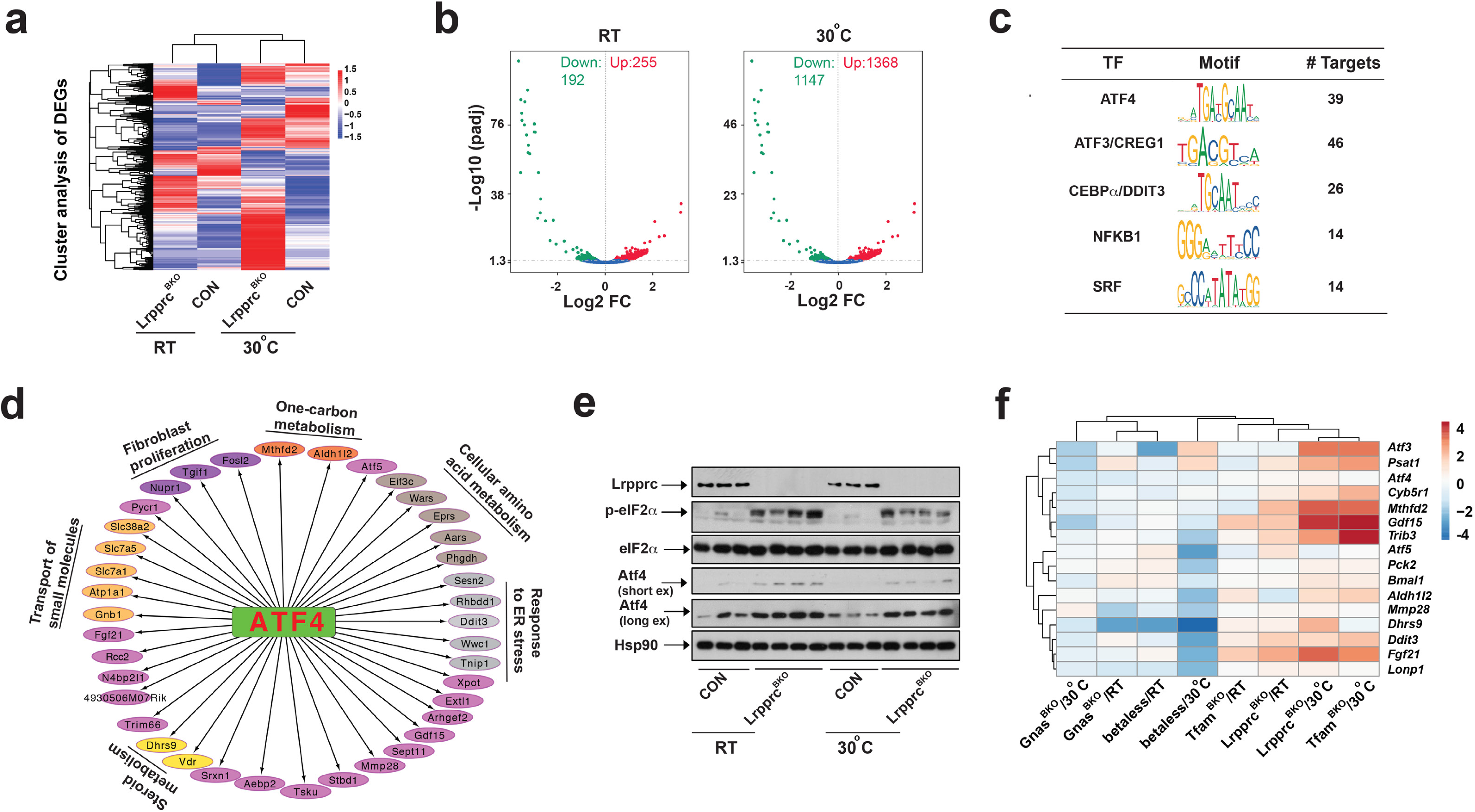
NOSEMPE specifically activates ATF4-ISR in brown adipocytes. **(a)** Heatmap showing hcluster analysis of DEGs in the BAT of male CON and Lrpprc^BKO^ mice at RT and 30°C. **(b)** Volcano plots showing significantly (p<0.05) down- or up-regulated genes in the BAT of Lrpprc^BKO^ mice at RT and 30°C. **(c)** List of enriched Transcript factors (TF) in commonly up-regulated DEGs in the BAT of CON and Lrpprc^BKO^ mice at RT and 30°C. Name, motif sequence and number of targets of each TF shown. **(d)** ATF4 signaling network. GO terms of ATF4 targets shown. **(e)** Immunoblots showing amounts of Lrpprc, p-eIF2α, total eIF2α, Atf4 and Hsp90 in the BAT of ∼8-12-week-old male CON and Lrpprc^BKO^ mice at normal chow at both RT and 30°C. **(f)** Clustering analysis of log2 fold changes of known ATF4 target genes in the BAT of mouse models with defective thermogenesis (Lrpprc^BKO^, Tfam^BKO^, Gnas^BKO^ and betaless mice) at normal chow at both RT and 30°C.

On the other hand, pathways of immune cell activation and protein homeostasis were highly enriched among the upregulated 165 genes at both RT and 30°C (**Supplementary Fig.5d, e**). Cis-regulatory sequence analysis using iRegulon predicted that ATF4 and its downstream transcription factors ATF3 and DDIT3 were the top regulators of the upregulated DEGs (**Fig2.c**). For example, 39 (out of 165) genes were predicted to contain putative ATF4 response element “TTGCATCA” on their promoter regions. These genes regulated diverse cellular processes, such as response to ER stress, cellular amino acid metabolism, transport of small molecules, one-carbon metabolism, steroid metabolism and fibroblast proliferation (**Fig.2d**). We experimentally confirmed that the mRNA levels of the ATF4 targets were upregulated in the BAT of Lrpprc^BKO^ mice at both RT and 30°C fed with either normal chow or HFD (**Supplementary Fig.6a**). Interesting, several ATF4 targets, such as Cyb5r1, Pck2 and Lonp1, were also identified amongst the upregulated proteins in the BAT mitochondria from both Tfam^BKO^ and Lrpprc^BKO^ mice (**Supplementary Fig.2i**). Thus, this transcriptomic profiling study indicates that ATF4 transcription network, also called the integrated stress response (ISR), is activated by NOSEMPE in brown adipocytes.

The ISR is centrally controlled by the phosphorylation of eukaryotic translation initiation factor eIF2α. When phosphorylated, it specifically induces ATF4 translation and its target gene expression ^27^. Indeed, we observed phosphor-eIF2α and Atf4 protein were elevated, along with known ATF4 targets in the BAT of Lrpprc^BKO^ mice at RT and 30°C (**Fig.2e**). Tfam^BKO^ mice, but not betaless mice, showed a similar induction of p-eIF2α in the BAT (**Supplementary Fig.6b, c**). Additional clustering analysis using these ATF4 targets in various mouse models with defective thermogenesis such as betaless, Gnas^BKO^, Lrpprc^BKO^, Tfam^BKO^ mice ^15, 24^ clearly demonstrated that ATF4-dependent ISR was specifically induced by NOSEMPE, but not a nonspecific adaptive response to defective thermogenesis in brown adipocytes (**Fig.2f**).

### Nonmitochondrial thermogenesis induced by NOSEMPE and ATF4 is dependent on proteome turnover

Since elevated ATF4 expression is sufficient for transcriptional induction of its targets *in vitro* and *in vivo* ^28, 29^, we then decided to ectopically overexpress ATF4 in brown adipocytes to examine its consequences on mitochondrial thermogenesis and nonmitochondrial thermogenesis. We have crossed ROSA-LSL-FlaghATF4 (Flag-tagged human ATF4 flanked by stop cassette in ROSA locus) with Ucp1-Cre mice, to generate brown adipocyte-specific ATF4 overexpressing mice (ATF4^BOX^) and their controls (**Supplementary Fig.7a**). Flag immunoblot confirmed the presence of Flag-ATF4 protein in the BAT of the ∼8-week-old ATF4^BOX^ mice at normal chow at RT (**Supplementary Fig.7b**). Similar to Lrpprc^BKO^ and Tfam^BKO^ mice, ATF4^BOX^ mice exhibited marked upregulation of ATF4 target gene network in the BAT at both RT and 30°C (**Fig.3a**), further confirming the ISR activation by ATF4 overexpression in the BAT. Thus, the ATF4^BOX^ mice phenocopied the Lrpprc^BKO^ and Tfam^BKO^ mice in regard to the brown adipocyte ISR activation specifically.

**Figure 3.**
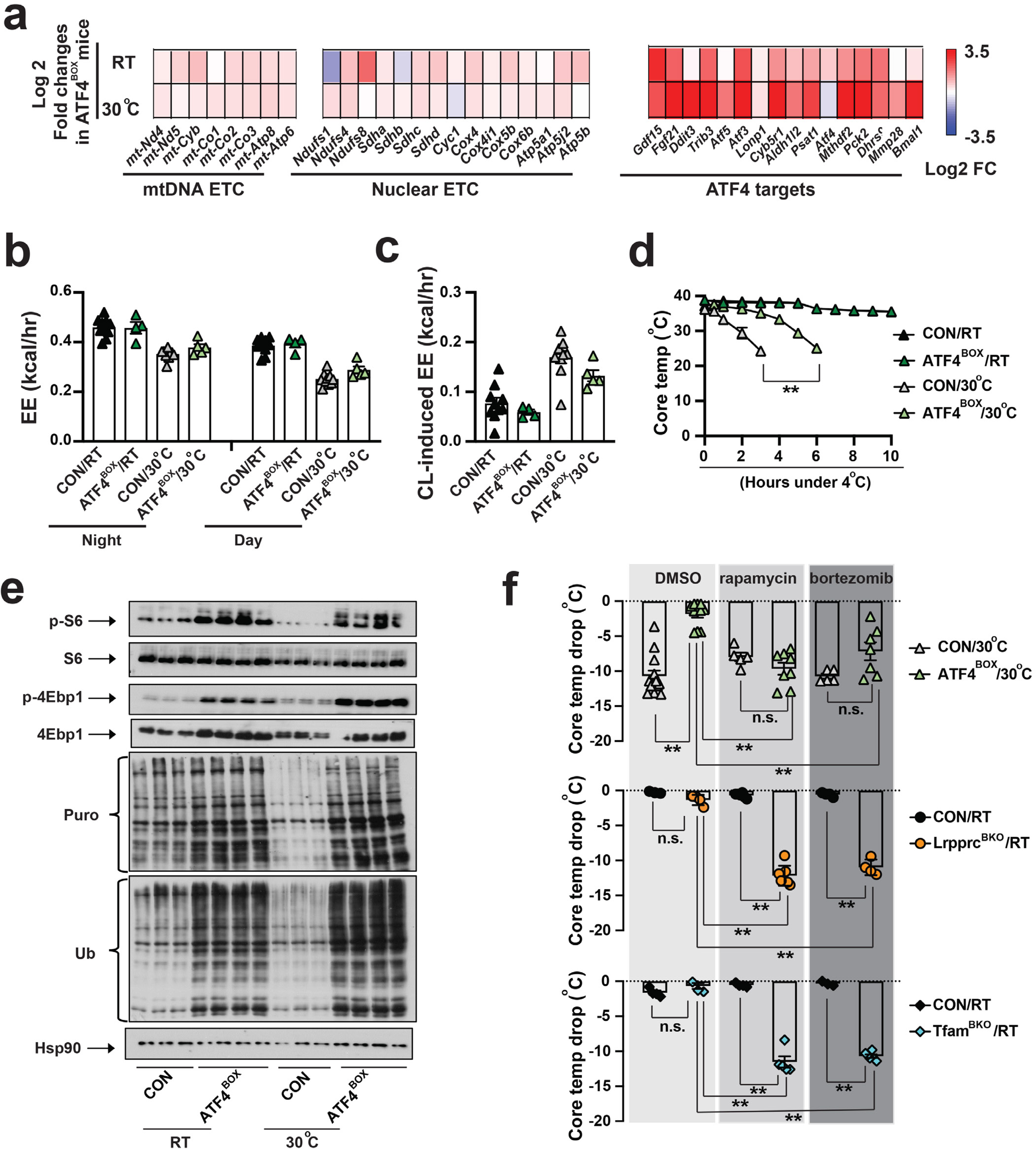
Brown adipocyte-specific ATF4 overexpression is sufficient to induce the nonmitochondrial thermogenesis fueled by proteome turnover. **(a)** Heatmap showing the log2 fold changes of mtDNA-encoded and nuclear-encoded ETC subunits and ATF4 targets in the BAT from ∼8-12-week of male CON and brown adipocyte-specific ATF4 overexpression (ATF4^BOX^) mice housed at RT and 30°C. Sample size: CON/RT (n=8), ATF4^BOX^/RT (n=8), CON/30°C (n=4) and ATF4^BOX^/30°C (n=6). Average night and day EE **(b)** and hourly CL-induced EE **(c)** in ∼8-12-week old male CON and ATF4^BOX^ mice for three days at RT and 30°C. Sample size: CON/RT (n=10), ATF4^BOX^/RT (n=4), CON/30°C (n=10) and ATF4^BOX^/30°C (n=5). **(d)** CTT of ∼8-12-week old male and female CON and ATF4^BOX^ mice housed at RT and 30°C. Sample size: CON/RT (n=5), ATF4^BOX^/RT (n=5), CON/30°C (n=6) and ATF4^BOX^/30°C (n=7). **(e)** Immunoblots of p-S6, total S6, p-4Ebp1, total 4Ebp1, puromycylated protein, ubiquitinated protein and Hsp90 in the BAT of ∼10-week-old male CON and ATF4^BOX^ mice housed at RT and 30°C. **(f)** Core temperature drop of ∼10-week-old male and female CON and ATF4^BOX^ mice with pretreatment of DMSO or 4mg/kg rapamycin, or 0.625mg/kg bortezomib after 3 hours 4°C CTT from 30°C. Core temperature drop of ∼10-week-old male and female Lrprp^BKO^ and Tfam^BKO^ mice and their relative controls with pretreatment of DMSO or rapamycin or bortezomib after 8 hours 4°C CTT from RT. Sample size: For ATF4^BOX^mice: CON/30°C/DMSO (n=13), ATF4^BOX^/30°C/DMSO (n=11), CON/30°C/rapamycin (n=5), ATF4^BOX^/30°C/rapamycin (n=8), CON/30°C/bortezomib (n=5) and ATF4^BOX^/30°C/ bortezomib (n=7). For Lrpprc^BKO^ mice: CON/RT/DMSO (n=4), Lrpprc^BKO^/RT/DMSO (n=4), CON/RT/rapamycin (n=6), Lrpprc^BKO^/RT/rapamycin (n=7), CON/RT/bortezomib (n=4) and Lrpprc^BKO^/RT/bortezomib (n=4). For Tfam^BKO^ mice: CON/RT/DMSO (n=4), Tfam^BKO^/RT/DMSO (n=4), CON/RT/rapamycin (n=4), Tfam^BKO^/RT/rapamycin (n=5), CON/RT/bortezomib (n=3) and Tfam^BKO^/RT/bortezomib (n=6). Data was presented as average ± SEM. Student t-test. n.s.: non-significant; **: p<0.01.

Indirect calorimetry experiments showed that the ATF4^BOX^ mice exhibited the similar βAR-induced adaptive thermogenesis as control mice at both ambient temperatures, so did the basal EE, RER, food intake and physical activity (**Fig.3b, c, Supplementary Fig.7d-f**). This was consistent with the lack of changes in BAT thermogenic and ETC (nuclear- and mtDNA-encoded) gene expression (**Fig.3a, Supplementary Fig.7c**). Therefore, ATF4 overexpression in brown adipocytes does not affect mitochondrial ETC gene expression and βAR-induced adaptive thermogenesis. When housed at RT, ATF4^BOX^ mice were cold resistant as the control mice during CTT, because mitochondrial thermogenesis was not affected by the ATF4 overexpression in brown adipocytes (**Fig.3d**). Thermoneutral housing (lack of sympathetic inputs) reduces PGC1α expression and mitochondrial biogenesis in brown adipocytes, similar to Gnas deficiency ^24^. Therefore, wild-type mice, acclimated to thermoneutrality, rapidly drop their core temperature upon CTT due to diminished mitochondrial thermogenesis. However, ATF4^BOX^ mice exhibited enhanced cold resistance during CTT from 30°C (**Fig.3d**), further suggesting the presence of ATF4-dependent nonmitochondrial thermogenesis in brown adipocytes.

Besides Ucp1-mediated uncoupling in mitochondria, heat can be generated through futile cycles where two metabolic pathways operate simultaneously in opposite directions ^30, 31^, such as calcium cycle, creatine cycle, triglyceride-fatty acid cycle, and glycolysis-gluconeogenesis cycle ^32–35^. Proteome turnover, the coupled protein synthesis and degradation could be a potential thermogenic mechanism especially in brown adipocytes, by wasting ATP to generate heat as the byproduct ^36^. As numerous ATF4 signature genes, involved multiple steps in amino acid synthesis (*Prcy1, Phgdh, Psat1* and *Mthdf2*), amino acid transporters (*Slc7a1, Slc7a5, Slc38a2*), and protein synthesis (aminoacyl-tRNA synthetases: *Wars, Eprs, Aars*) were upregulated by NOSEMPE and ATF4 overexpression in brown adipocytes (**Fig.2d**), we reason that ATF4 may promote proteome turnover primarily due to its anabolic action on protein synthesis.

We evaluated rates of protein synthesis in the BAT of ATF4^BOX^ mice at both RT and 30°C. First, ribosome protein S6 and mRNA translation repressor 4Ebp1 were highly phosphorylated in the BAT of ATF4^BOX^ mice (**Fig.3e**), reflecting a higher activity of mTORC1 (the master regulator of protein synthesis) ^37^. We then evaluated the global protein synthesis rate by puromycin labeling ^38^. Indeed, puromycylated proteins were elevated in the BAT of ATF4^BOX^ mice (**Fig.3e**). Next, a protein synthesis inhibitor (rapamycin) was used to determine the contributions of protein synthesis to the increased cold resistance phenotype of the ATF4^BOX^ mice acclimated at 30°C. Pretreatment of 4mg/kg rapamycin fully abolished elevated p-S6, p-4Ebp1, total puromycylated proteins in the BAT of ATF4^BOX^ mice (**Supplementary Fig.8a**). Importantly, ATF4^BOX^ mice were no longer cold resistant after rapamycin treatment, although control mice behaved similarly during CTT with or without rapamycin (**Fig.3f**). In contrast, global protein synthesis was not affected in the muscle of the ATF4^BOX^ mice, and rapamycin did not affect the phosphorylation of S6 and 4Ebp1, and the rates of global protein synthesis in the muscle (**Supplementary Fig.8b**), suggesting the effect of rapamycin in cold resistance phenotype in the ATF4^BOX^ mice was due to inhibition of mTORC1-dependent protein synthesis specifically in the BAT.

In parallel, the global ubiquitinated proteins were also elevated (**Fig.3e**) and proteasome inhibitor (bortezomib) had a similar effect in ATF4^BOX^ mice during CTT (**Fig.3f**). Inhibition of protein synthesis by rapamycin also reduced total ubiquitinated proteins in the BAT of ATF4^BOX^ mice (**Supplementary Fig.8a**), suggesting that protein synthesis and degradation were coupled and the accelerated proteome turnover in brown adipocytes contributed to the cold resistance phenotype in the ATF4^BOX^ mice.

We next assessed BAT proteome turnover in Lrpprc^BKO^ mice similarly. Phosphorylated S6 and 4Ebp1 and puromycylated and ubiquitinated proteins were elevated in the BAT of Lrpprc^BKO^ mice (**Supplementary Fig.9a**). Treatment with rapamycin or bortezomib suppressed the cold resistance phenotype of the Lrpprc^BKO^ mice without any noticeable changes in control mice in CTT (**Fig.3f**). Tfam^BKO^ mice, another mouse model of BAT NOSEMPE, showed similar phenotypes (**Fig.3f, Supplementary Fig.9b**). Collectively, ATF4-mediated proteome turnover constitutes a mechanism of nonmitochondrial thermogenesis in brown adipocytes.

### NOSEMPE and ATF4 activation in brown adipocytes promote systemic metabolism

Next, we characterized the metabolic consequences of elevated nonmitochondrial thermogenesis by either BAT NOSEMPE or ATF4 overexpression. We found that male Lrpprc^BKO^ mice gained less body weight under normal chow feeding at both ambient temperatures (**Supplementary Fig.10a**). At ∼8-month of age, Lrpprc^BKO^ mice exhibited reduced adipose mass in BAT, subcutaneous inguinal WAT (iWAT) and visceral epididymal WAT (eWAT) (**Supplementary Fig.10b, c**).

Male Lrpprc^BKO^ mice gained less body weight under high fat diet (HFD) (**Fig.4a**). The difference in body weight was already evident after 4 weeks on HFD and became greater after longer-period HFD (**Fig.4a, c**). After 12-week HFD, body weight gains of Lrpprc^BKO^ mice were only 28% and 37% of those of control littermates. The body weight difference was mainly contributed by reduction of fat mass. Fat percentage was increased in control mice by four-fold by 12-week HFD, but it barely increased in Lrpprc^BKO^ mice (**Fig.4d**). Fat depots like iWAT and eWAT showed progressively reduced weight under HFD and contained smaller adipocytes, and BAT only showed reduced weight after 12-week HFD (**Fig.4b, c**). The adipocyte size in eWAT was also reduced in Lrpprc^BKO^ mice after HFD (**Fig.4e, f**).

**Figure 4.**
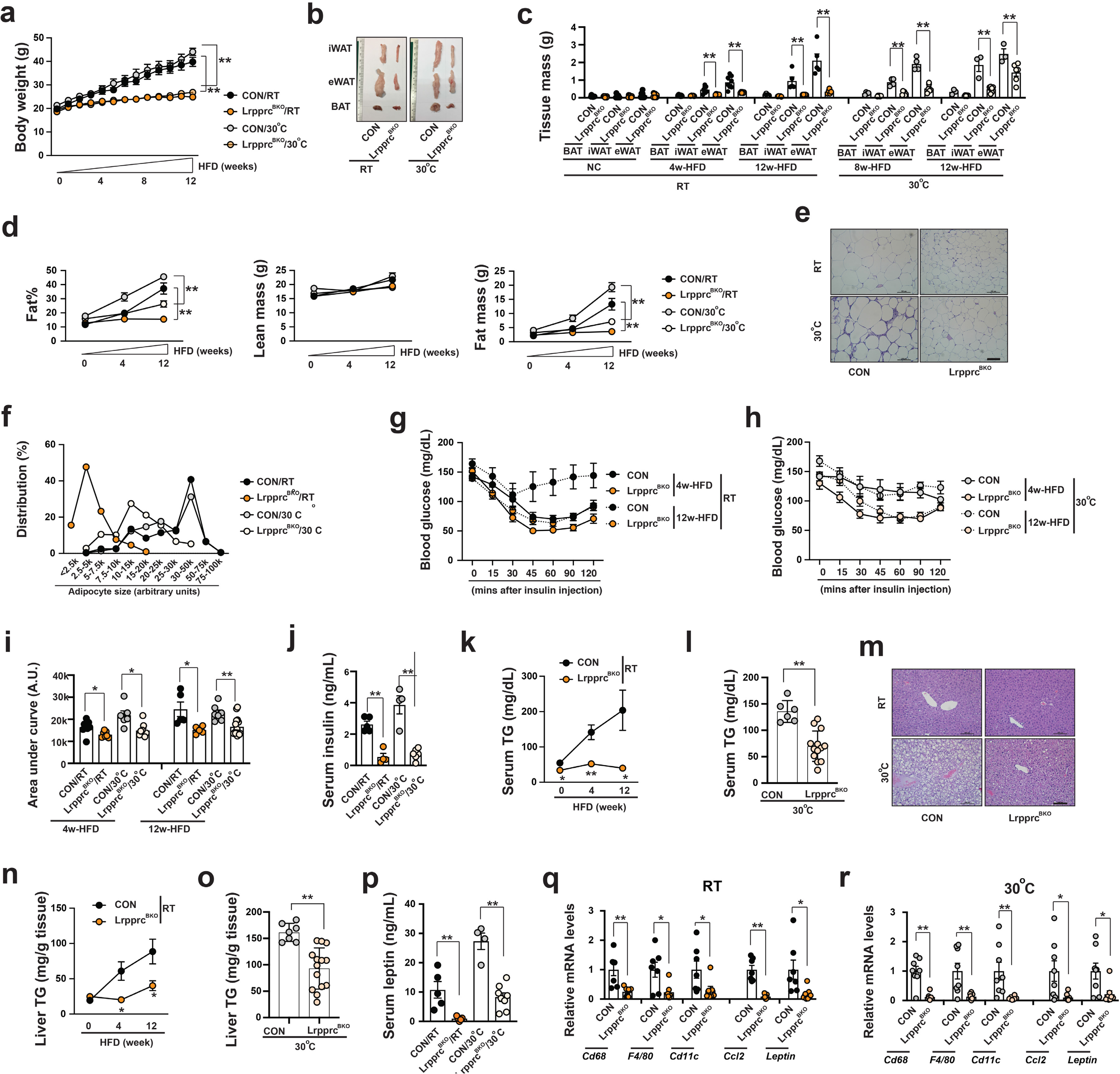
Lrpprc^BKO^ mice exhibit improved systemic metabolism under HFD. **(a)** Body weight of male CON and Lrpprc^BKO^ mice under 12-week HFD at RT and 30°C. Sample size: CON/RT (n=12), Lrpprc^BKO^/RT (n=11), CON/30°C (n=7) and Lrpprc^BKO^/30°C (n=15). **(b)** Representative images of dissected iWAT, eWAT and BAT from male CON and Lrpprc^BKO^ mice after 12-week HFD. **(c)** Tissue mass of eWAT, iWAT, and BAT of male CON and Lrpprc^BKO^ mice at normal chow (NC) and after HFD. Sample size: male CON/NC/RT (n=8), Lrpprc^BKO^/NC/RT (n=10), CON/4w-HFD/RT (n=8), Lrpprc^BKO^/4w-HFD/RT (n=6), CON/12w-HFD/RT (n=5), Lrpprc^BKO^/12w-HFD/RT (n=5), CON/8w-HFD/30°C (n=4), Lrpprc^BKO^/8w-HFD/30°C (n=6), CON/12w-HFD/30°C (n=6) and Lrpprc^BKO^/12w-HFD/30°C (n=13). **(d)** Fat percentage, lean and fat mass of male CON and Lrpprc^BKO^ mice before and after 4-week and 12-week HFD at RT and 30°C. Sample size: CON/NC/RT (n=3), Lrpprc^BKO^/NC/RT (n=7), CON/4w-HFD/RT (n=10), Lrpprc^BKO^/4w-HFD/RT (n=9), CON/12w-HFD/RT (n56), Lrpprc^BKO^/12w-HFD/RT (n=5), CON/NC/30°C (n=7), Lrpprc^BKO^/NC/30°C (n=6), CON/4w-HFD/30°C (n=6), Lrpprc^BKO^/4w-HFD/30°C (n=8), CON/12w-HFD/30°C (n=6) and Lrpprc^BKO^/12w-HFD/30°C (n=13). **(e)** Representative H&E staining of eWAT from male CON and Lrpprc^BKO^ mice after 12-week HFD. Scale bar: 100 μm. **(f)** Adipocyte size distribution in eWAT from male CON and Lrpprc^BKO^ mice after 12-week HFD. Total adipocytes counted: CON/RT (n=200), Lrpprc^BKO^/RT (n=519), CON/30°C (n=347) and Lrpprc^BKO^/30°C (n=666). Serum glucose levels during ITT in male CON and Lrpprc^BKO^ mice after 4-week and 12-week HFD at RT **(g)** and 30°C **(h)**. Sample size: CON/4w-HFD/RT (n=8), Lrpprc^BKO^/4w-HFD/RT (n=7), CON/12w-HFD/RT (n=5), Lrpprc^BKO^/12w-HFD/RT (n=5), CON/4w-HFD/30°C (n=6), Lrpprc^BKO^/4w-HFD/30°C (n=9), CON/12w-HFD/30°C (n=7) and Lrpprc^BKO^/12w-HFD/30°C (n=15). **(i)** Area under the curve (AUC) values of glucose levels in ITTs showed. **(j)** Serum insulin levels in male CON and Lrpprc^BKO^ mice after 12-week HFD. Sample size: CON/RT (n=5), Lrpprc^BKO^/RT (n=4), CON/30°C (n=4), Lrpprc^BKO^/30°C (n=6). **(k)** Serum triglyceride contents of male CON and Lrpprc^BKO^ mice after HFD at RT. Sample size: Sample size: CON/NC/RT (n=6), Lrpprc^BKO^/NC/RT (n=10), CON/4w-HFD/RT (n=8), Lrpprc^BKO^/4w-HFD/RT (n=10), CON/12w-HFD/RT (n=5) and Lrpprc^BKO^/12w-HFD/RT (n=5). **(l)** Serum triglyceride contents of male CON and Lrpprc^BKO^ mice after 12-week HFD at 30°C. Sample size: CON/12w-HFD/30°C (n=6) and Lrpprc^BKO^/12w-HFD/30°C (n=13). **(m)** Representative H&E staining of liver from male CON and Lrpprc^BKO^ mice after 12-week HFD. Scale bar: 100 μm. **(n)** Liver triglyceride contents of male CON and Lrpprc^BKO^ mice after HFD at RT. Sample size: CON/NC/RT (n=6), Lrpprc^BKO^/NC/RT (n=7), CON/4w-HFD/RT (n=7), Lrpprc^BKO^/4w-HFD/RT (n=6), CON/12w-HFD/RT (n=5) and Lrpprc^BKO^/12w-HFD/RT (n=5). **(o)** Liver triglyceride contents of male CON and Lrpprc^BKO^ mice after 12-week HFD at 30°C. Sample size: CON/12w-HFD/30°C (n=7) and Lrpprc^BKO^/12w-HFD/30°C (n=14). **(p)** Serum leptin levels in male CON and Lrpprc^BKO^ mice after 12-week. Sample size: CON/RT (n=5), Lrpprc^BKO^/RT (n=5), CON/30°C (n=4) and Lrpprc^BKO^/30°C (n=8). Q-PCR analysis of mRNA levels of macrophage markers (*Cd68, F4/80* and *Cd11c*) and pro-inflammatory cytokines (*Ccl2* and *Leptin*) in eWAT of male CON and Lrpprc^BKO^ mice after 12-week HFD at RT **(q)** and 30°C **(r)**. Sample size: CON/RT (n=7), Lrpprc^BKO^/RT (n=7), CON/30°C (n=8) and Lrpprc^BKO^/30°C (n=8). Data was presented as average ± SEM. Student t-test. *: p<0.05 and **: p<0.01.

We also measured other HFD-induced metabolic parameters in male Lrpprc^BKO^ mice. Systemic insulin sensitivity was improved in Lrpprc^BKO^ mice (**Fig.4g-i**) as early as after 4-week HFD, and serum insulin levels were significantly reduced at 12-week HFD (**Fig.4j**). HFD-induced hypertriglyceridemia was completely inhibited in Lrpprc^BKO^ mice. Serum triglyceride contents were elevated by about four-fold (over 200 mg/dL) in control mice at RT but remained at lower levels (<40 mg/dL) in Lrpprc^BKO^ mice after 12-week HFD (**Fig.4k**). Similar results were also obtained at 30°C (**Fig.4l**). HFD also induced ectopic triglyceride accumulation in the liver in control mice, which was also absent in Lrpprc^BKO^ mice (**Fig.4m-o**). HFD-induced adipose inflammation was suppressed by Lrpprc deficiency in brown adipocytes. Q-PCR analysis showed macrophage markers (*Cd68, F4/80* and *Cd11c*) and pro-inflammatory cytokines (*Ccl2* and *Leptin*) were reduced in the eWAT of Lrpprc^BKO^ mice after 12-week HFD (**Fig.4q-r**) and serum leptin levels were also reduced (**Fig.4p**). Thus, Lrpprc deficiency in brown adipocytes in BAT leads to the protection against HFD-induced obesity, insulin resistance, hepatic steatosis, hypertriglyceridemia, and adipose inflammation, despite defective thermogenesis in BAT. The reduced adiposity and liver TG phenotypes were also observed in female Lrpprc^BKO^ mice (**Supplementary Fig.11a-d**). Thus, BAT NOSEMPE improves systemic metabolism in the Lrpprc^BKO^ and Tfam^BKO^ mice despite a complete absence of βAR-induced adaptive thermogenesis.

In order to determine whether ATF4 activation in wild-type brown adipocytes is sufficient to drive systemic metabolic benefits, we performed HFD experiments similarly on ATF4^BOX^ mice at both RT and 30°C. Indeed, ATF4-ISR remained elevated in the BAT of ATF4^BOX^ mice even after HFD (**Fig.5b**). ATF4^BOX^ mice gained less body weight, which was contributed by reduced adiposity (**Fig.5a, c, d**). Again, other metabolic parameters such as increased adipocyte size and inflammation in white adipose tissue (**Fig.5e, f, m**), systemic insulin resistance (**Fig.5g-i**), hepatosteatosis and hyperlipidemia (**Fig.5j-l**) were all suppressed by ATF4 overexpression in brown adipocytes. Taken together, ATF4 activation in brown adipocytes is sufficient to induce nonmitochondrial thermogenesis from BAT (without an increase of βAR-induced adaptive thermogenesis) and improves systemic metabolism.

**Figure 5.**
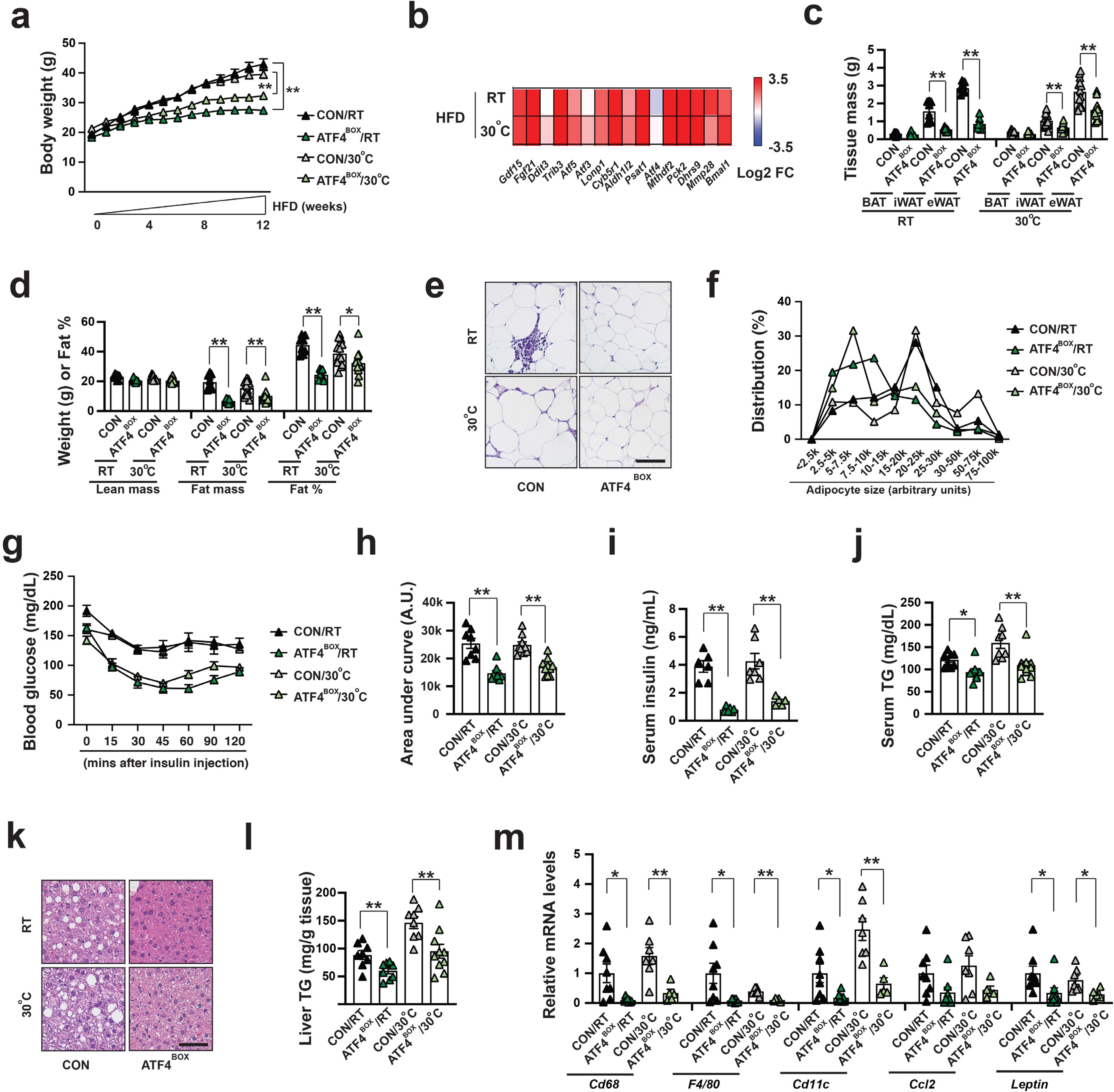
ATF4 activation in brown adipocytes is sufficient to drive metabolic fitness. **(a)** Body weight of male CON and ATF4^BOX^ mice after 12-week HFD at RT and 30°C. Sample size: CON/RT (n=8), ATF4^BOX^/RT (n=8), CON/30°C (n=17) and ATF4^BOX^/30°C (n=18). **(b)** Heatmap showing log2 fold changes of known ATF4 target genes in the BAT of ATF4^BOX^ mice after 4-week HFD at both RT and 30°C. Sample size: CON/RT (n=8), ATF4^BOX^/RT (n=7), CON/30°C (n=4) and ATF4^BOX^/30°C (n=6). **(c)** Tissue mass of eWAT, iWAT, and BAT of male CON and ATF4^BOX^ mice after 12-week HFD. Sample size: male CON/RT (n=8), ATF4^BOX^/RT (n=8), CON/30°C (n=13) and ATF4^BOX^/30°C (n=13). **(d)** Lean mass, fat mass, and fat percentage of male CON and Lrpprc^BKO^ mice after 12-week HFD. Sample size: CON/RT (n=8), ATF4^BOX^/RT (n=8), CON/30°C (n=13) and ATF4^BOX^/30°C (n=13). **(e)** Representative H&E staining of eWAT from male CON and ATF4^BOX^ mice after 12-week HFD. Scale bar: 100 μm. **(f)** Adipocyte size distribution in eWAT from male CON and ATF4^BOX^ mice after 12-week HFD. Total adipocytes counted: CON/RT (n=2647), ATF4^BOX^/RT (n=2312), CON/30°C (n=1868) and ATF4^BOX^/30°C (n=2650). **(g)** Serum glucose levels during ITT in male CON and ATF4^BOX^ mice after 12-week HFD at RT and 30°C. **(h)** Area under the curve (AUC) values of glucose levels in ITTs showed. Sample size: CON/RT (n=8), ATF4^BOX^/RT (n=8), CON/30°C (n=9) and ATF4^BOX^/30°C (n=10). **(i)** Serum insulin levels in male CON and ATF4^BOX^ mice after 12-week HFD. Sample size: CON/RT (n=6), ATF4^BOX^/RT (n=6), CON/30°C (n=7), ATF4^BOX^/30°C (n=5). **(j)** Serum triglyceride contents of male CON and ATF4^BOX^ mice after 12-week HFD. Sample size: CON/RT (n=8), ATF4^BOX^/RT (n=8), CON/30°C (n=8) and ATF4^BOX^/30°C (n=10). **(k)** Representative H&E staining of liver from male CON and ATF4^BOX^ mice after 12-week HFD. Scale bar: 100 μm. **(l)** Liver triglyceride contents of male CON and ATF4^BOX^ mice after 12-week HFD. Sample size: CON/RT (n=8), ATF4^BOX^/RT (n=8), CON/30°C (n=8) and ATF4^BOX^/30°C (n=10). **(m)** Q-PCR analysis of mRNA levels of macrophage markers (*Cd68, F4/80* and *Cd11c*) and pro-inflammatory cytokines (*Ccl2* and *Leptin*) in eWAT of male CON and ATF4^BOX^ mice after 12-week HFD. Sample size: CON/RT (n=8), ATF4^BOX^/RT (n=8), CON/30°C (n=7) and ATF4^BOX^/30°C (n=5). Data was presented as average ± SEM. Student t-test. *: p<0.05 and **: p<0.01.

### ATF4 activation is required for NOSEMPE-induced nonmitochondrial thermogenesis and metabolic benefits

While Atf4 global knockout mice are lean ^39, 40^, Atf4’s specific role in brown adipocytes is unknown. To address this question, we have generated brown adipocyte-specific Atf4 knockout mice (*Ucp1-Cre;Atf4^f/f^*, Atf4^BKO^). Atf4 deficiency in brown adipocytes reduced ATF4 target gene expression but did not affect multilocular morphology at RT and mitochondrial ETC gene expression at RT and 30°C (**Supplementary Fig12.a, b**). Indirect calorimetry experiments showed that Atf4 deficiency did not affect basal and CL-induced EE, RER, food intake and physical activity (**Supplementary Fig12.c-g**). Most importantly, HFD-induced obesity, insulin resistance, hepatosteatosis and hyperlipidemia were not altered by Atf4 deficiency (**Supplementary Fig12.h-o**).

We then determined whether Atf4 activation in brown adipocytes was required for the nonmitochondrial thermogenesis and metabolic improvements in Lrpprc^BKO^ mice. We have generated brown adipocyte-specific Lrpprc and Atf4 double knockout mice (*Ucp1-Cre;Lrpprc^f/f^;Atf4^f/f^*, Lrpprc;Atf4^BKO^). Lrpprc and Atf4 double deficient brown adipocytes were unilocular and exhibited reduced mtDNA ETC gene expression and p-eIF2α, similar to the Lrpprc^BKO^ mice (**Supplementary Fig.13a, b, Fig.6a**). Consistently, the Lrpprc;Atf4^BKO^ mice at 30°C also lacked CL-induced EE due to the defect of mtDNA gene expression in brown adipocytes (**Fig.6b,c**), but without noticeable changes in basal EE, food intake and physical activity (**Supplementary Fig.14a-c**). However, the upregulation of ATF4 target genes and rates of global protein synthesis and degradation induced by Lrpprc deficiency was attenuated by additional Atf4 deficiency in brown adipocytes (**Supplementary Fig.13b, Fig.6a**). The Lrpprc;Atf4^BKO^ mice were no longer cold resistant in CTT, although Atf4 deficiency itself in brown adipocytes did not affect core body temperature during CTT (**Supplementary Fig.13c**). We then characterized the full spectrum of metabolic performance of Lrpprc;Atf4^BKO^ mice at 30°C. Compared to the Lrpprc^BKO^ mice, the Lrpprc;Atf4^BKO^ mice were no longer protected against HFD-induced obesity, adipocyte hypertrophy, systemic insulin resistance, hepatosteatosis and hyperlipidemia (**Fig.6d-h, Supplementary Fig.15a-f**). Thus, Atf4 activation is required for the NOSEMPE-induced, proteome turnover-fueled nonmitochondrial thermogenesis in brown adipocytes, and Atf4 deletion fully reverses metabolic benefits without affecting mitochondrial respiration and mitochondrial thermogenesis in the Lrpprc^BKO^ mice.

**Figure 6.**
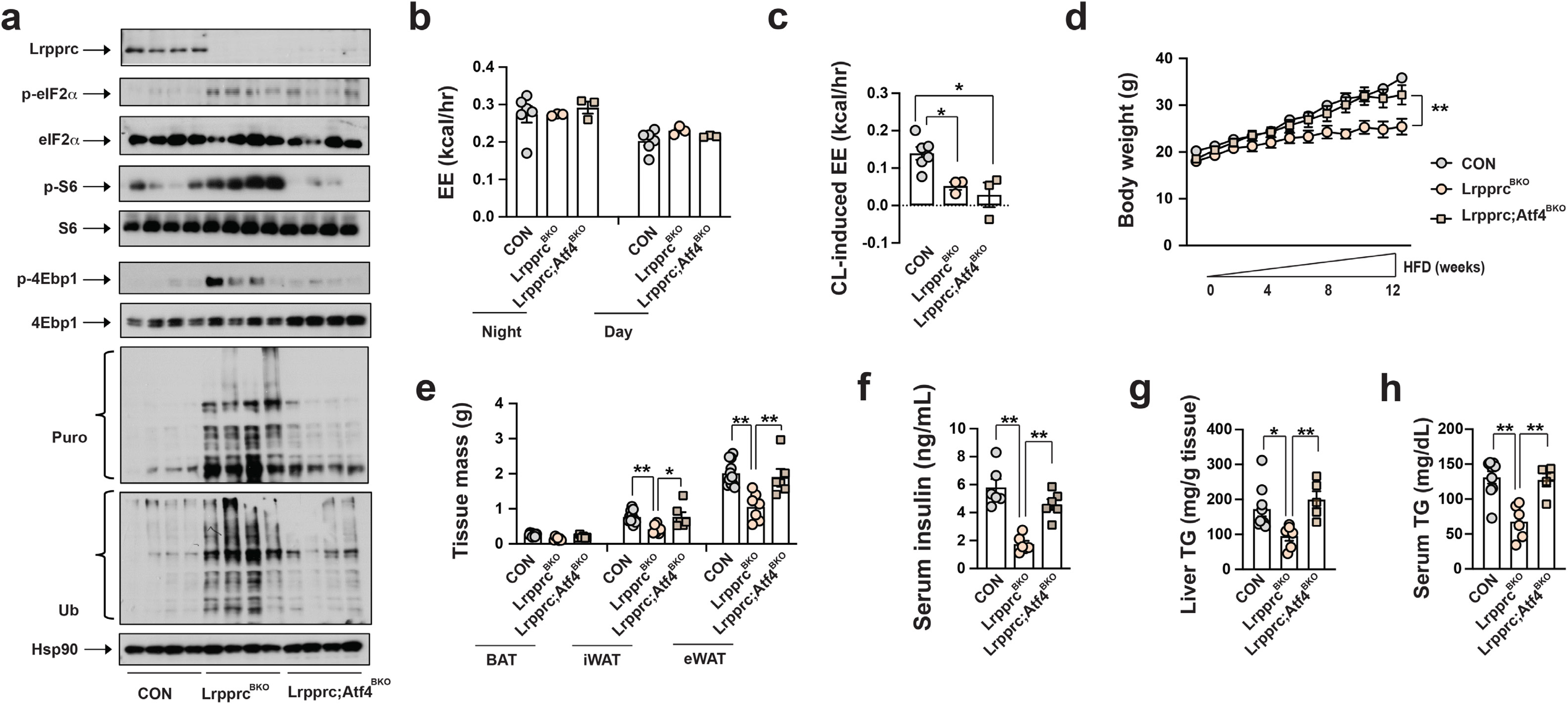
Atf4 activation is required for augmented nonmitochondrial thermogenesis and metabolic fitness in Lrpprc^BKO^ mice at 30°C. **(a)** Immunoblots of Lrpprc, p-eIF2α, total eIF2α, p-S6, total S6, p-4Ebp1, total 4Ebp1, puromycylated protein, ubiquitinated protein and Hsp90 in the BAT of ∼10-week-old male CON, Lrpprc^BKO^ and Lrpprc;Atf4^BKO^ mice housed at 30°C. Average night and day EE **(b)** and hourly CL-induced EE **(c)** in ∼10-week old male CON, Lrpprc^BKO^ and Lrpprc;Atf4^BKO^ mice for three days. Sample size: CON (n=6), Lrpprc^BKO^ (n=3), and Lrpprc;Atf4^BKO^ (n=3). **(d)** Body weight of male CON, Lrpprc^BKO^ and Lrpprc;Atf4^BKO^ mice under 12-week HFD at 30°C. **(e)** Tissue mass of eWAT, iWAT, and BAT of male CON, Lrpprc^BKO^ and Lrpprc;Atf4^BKO^ mice after 12-week HFD. Sample size: CON (n=11), Lrpprc^BKO^ (n=7), and Lrpprc;Atf4^BKO^ (n=6). Serum insulin **(f),** serum triglyceride contents and **(g)** and liver triglyceride contents **(h)** of male CON, Lrpprc^BKO^ and Lrpprc;Atf4^BKO^ mice after HFD. Sample size: CON (n=9), Lrpprc^BKO^ (n=6), and Lrpprc;Atf4^BKO^ (n=5). Data was presented as average ± SEM. Student t-test. *: p<0.05 and **: p<0.01.

## DISCUSSION

Comparative studies have suggested that the appearance of thermogenic BAT in endotherms provides an advantage to survive in cold environment throughout evolution ^41^. Mitochondrial respiration has been long regarded as the essential component of BAT adaptive thermogenesis. Thus, active brown adipocytes in rodents and humans are defined by three criteria: multilocular in morphology, rich in mitochondria and positive for UCP1. Divergent from this “mitochondrial thermogenesis” centric view, our studies demonstrate that the unilocular and mitochondrial respiration-defective brown adipocytes in Lrpprc^BKO^ and Tfam^BKO^ mice (especially at thermoneutrality) can also efficiently promote systemic metabolic health. Since adult humans mostly live at the thermoneutral condition, it is feasible that these adipocytes will have the potential to regulate systemic metabolism through the mechanism observed in mice with BAT NOSEMPE or ATF4 activation.

Moreover, a new concept termed as “nonmitochondrial thermogenesis” is emerging. Distinct from the well-studied mitochondrial thermogenesis (the βAR-dependent heat production inside mitochondria), this nonmitochondrial thermogenesis can operate without βAR stimulation. When housed at RT with mild cold stress, wild-type mice process activated mitochondrial thermogenesis and minimal nonmitochondrial thermogenesis. Gnas^BKO^ (or betaless or Ucp1 KO) mice or wild-type mice at thermoneutrality lose mitochondrial thermogenesis (without changes in nonmitochondrial thermogenesis), consequently showing defective thermoregulation in CTT. Even though Tfam^BKO^ and Lrpprc^BKO^ mice also lack mitochondrial thermogenesis completely at RT, they can still maintain their core temperature during CTT through augmented ATF4-dependent nonmitochondrial thermogenesis. Furthermore, ATF4^BOX^ mice after thermoneutral housing even exhibited enhanced cold resistance in CTT. Collectively, mitochondrial thermogenesis and nonmitochondrial thermogenesis are both crucial for organismal thermoregulation, and nonmitochondrial thermogenesis can even compensate for the deficiency of mitochondrial thermogenesis *in vivo*.

However, there are still several unanswered questions regarding this nonmitochondrial thermogenesis. Cellular proteome balance is precisely controlled by the capacity, velocity and fidelity of protein synthesis and degradation in the cytosol ^42^, and its maintenance is the most energy-consuming process (utilizing about one-third of total energy) ^43–45^. Although we have demonstrated the “NOSEMPE→ATF4→proteome turnover” pathway as a candidate mechanism for nonmitochondrial thermogenesis in brown adipocytes, its molecular details are not fully unveiled yet. For example, whether the turnover rates of specific classes of proteins or global proteome are accelerated in response to NOSEMPE and/or ATF4 activation is undetermined yet ^46, 47^. The protein synthesis-promoting effect of ATF4 has been observed in other cell types *in vitro* ^48–51^ and *in vivo* ^52^. How this increased protein synthesis is coupled with increased degradation in brown adipocytes remains unknown, although the ribosome-associated protein quality control ^53–55^ may represent one possible mechanism.

The relative contributions of mitochondrial thermogenesis and nonmitochondrial thermogenesis in brown adipocytes to systemic metabolism vary. For example, brown adipocyte-specific Irf4 overexpression mice (Irf4^BOX^) did show enhanced mitochondrial thermogenesis in brown adipocytes and a slight reduction in HFD-induced obesity, due to increased mitochondrial biogenesis ^10^. While Gnas^BKO^ and brown adipocyte-specific Hdac3 knockout mice (Hdac3^BKO^) lacked the mitochondrial thermogenesis but gained similar body weight under HFD ^24, 56^. We have demonstrated that mice with elevated nonmitochondrial thermogenesis in brown adipocytes (Lrpprc^BKO^, Tfam^BKO^, and ATF4^BOX^ mice), regardless of their activity in mitochondrial thermogenesis, exhibited most profound anti-obesity phenotype in reported mouse models using *Ucp1* promoter-driven constitutively active or inducible Cre ^10, 57, 58^. And removing the ATF4-dependent nonmitochondrial thermogenesis (without affecting mitochondrial thermogenesis) in Lrpprc^BKO^ mice reversed their metabolic benefits completely, suggesting a correlation between the nonmitochondrial thermogenesis in brown adipocytes and systemic metabolism. Energy wasting itself or byproducts of proteome turnover during this nonmitochondrial thermogenesis in brown adipocytes may potentially contribute to the metabolic benefits in Lrpprc^BKO^, Tfam^BKO^, and ATF4^BOX^ mice.

Future investigations into mechanisms and impacts of this novel nonmitochondrial thermogenesis will establish new paradigms for brown adipocyte biology beyond mitochondrial thermogenesis ^59^. Physiological and/or pathological conditions that affect any step of mtDNA-encoded protein expression, such as mitochondrial DNA maintenance, replication, RNA processing/maturation, ribosome assembly or protein translation ^60^, can potentially regulate systemic metabolism by inducing NOSEMPE in brown adipocytes. Mouse models with BAT NOSEMPE described here can be exploited to develop new nuclear imaging techniques to visualize the nonmitochondrial thermogenesis in brown adipocytes, which may be potentially employed in clinical practice in humans. Finally, targeting ATF4-dependent proteome turnover in brown adipocytes, besides increasing βAR-induced adaptive thermogenesis to increase total energy expenditure, may represent novel therapeutic approaches to treat obesity and associated metabolic disorders.

## Methods

### Mouse models

ROSA-LSL-FlaghATF4 mice (#029394) ^29^ were obtained from JAX. Lrpprc^f/f^, Atf4^f/f^, Ucp1-Cre (JAX #024670), and betaless mice were kindly provided by Drs. Nils-Göran Larsson, Christopher Adams, Evan Rosen, and Shingo Kajimura. Tfam^BKO^ and Gnas^BKO^ mice were characterized before ^15, 24^. Mice were housed in a temperature-controlled environment at 22°C under a 12h light:dark cycle with free access to water and food (PicoLab® Rodent Diet 20, #5053). For thermoneutral experiments, ∼4-week-old mice were placed in a 30°C rodent chamber (Power Scientific RIS52SD Rodent Incubator) for an additional 3-4 weeks to reach their thermoneutral zone. There were no inclusion/exclusion criteria for mice studies. Mice were in C57BL/6J background (except for Gnas^BKO^ mice). All animal experiments were approved by the UCSF Institutional Animal Care and Use Committee in adherence to US National Institutes of Health guidelines and policies.

### Metabolic studies

About 8-week-old mice were transferred to a 60% fat diet (Research Diets, D12492) housed at RT or 30°C. Body weight was monitored once a week. EchoMRI was performed following manufacturer’s instructions. For insulin tolerance test (ITT), mice were fasted 4-6 hours before intraperitoneal administration of insulin (Humulin; 0.75U kg^-1^). Blood glucose was measured from tail vein at indicated time points with a glucometer (Contour, Bayer). Serum and liver TG contents were measured by Infinity Triglycerides Reagents (Thermo Scientific, #TR22421). Serum insulin and leptin levels were measured by commercial ELISA kits (Alpco, #80-INSMSV-E01; Crystal Chem Inc, #90030).

### Indirect calorimetry measurements

Basal energy expenditure (EE) and CL-induced EE were calculated per mouse ^61, 62^. Investigators were blinded to the mouse genotypes for CLAMS, which was performed by the UCSF Diabetes and Endocrinology Research Center Metabolic Research Unit.

### ^18^F-fluorodeoxyglucose (FDG) uptake

A dedicated small animal PET/CT (Inveon, Siemens Medical Solutions, Malvern, PA) was used for all imaging procedures at room temperature. For consistent data acquisition, all animals were fasted overnight, at least 12 hours, before administration of ^18^F-fluorodeoxyglucose (FDG). FDG (3.94±0.17 MBq, range: 3.67-4.17 MBq) was administered intravenously via tail vein under anesthesia (2-2.5% isoflurane). Uptake time of 55 min (±1 min) was strictly followed before the start of the scan. During the uptake time, the animals were awake and kept warm over a temperature-controlled heating pad at 37 °C. Ten-minute static PET data were acquired for all animals, followed by CT under isoflurane (2-2.5%) anesthesia. The total imaging time was under 20 minutes. Once the data for PET and CT were acquired, reconstructions were performed using vendor-provided software. An iterative reconstruction algorithm with CT-based attenuation correction was used for PET, and a Feldkamp reconstruction algorithm modified for conebeam was used for CT. The reconstructed volumes were 128×128×159 matrices with a voxel size of 0.776383 mm × 0.776383 mm × 0.796 mm for PET, and 512×512×700 matrices with an isotropic voxel size of 0.196797 mm × 0.196797 mm × 0.196797 mm for CT. The CT acquisition parameters were: continuous 120 rotation steps over 220°, 80 kVp/500 μA tube voltage/current, and 175 ms exposure per step. Spherical VOIs (2 mm diameter) were drawn completely within brown adipose tissue, back of the cervical spine of each animal, and % injected dose per unit volume (%ID/ml) was calculated for analysis.

### Cold tolerance test (CTT)

∼8-week old male and female mouse was singly housed with free-access to food and water during CTT. The core body temperatures prior to and during 4°C cold exposure (at one-hour interval) were measured using BAT-12 Microprobe Thermometer with probe RET-3 (Physitemp). 4mg kg^-1^ rapamycin (TCI America, #TCR0097) or 0.625 mg kg^-1^ bortezomib (Selleck, #S1013) or DMSO (Sigma, #D8418) was injected intraperitoneally 1 hour prior to CTT.

### ETC Complex Activities

Frozen BAT tissue from about 8-week-old male and female mice was homogenized in 250 μL homogenization buffer (120 mM KCl, 20mM HEPES, 1mM EGTA, pH 7.4) by sonication (5 second pulse × 5, 60% power) using a Microson XL2000 Ultrasonic Cell Disruptor (Misonix). Protein was quantitated using the Bradford assay and all samples were diluted to a final concentration of 1μg/μl of protein. The spectrophotometric kinetic assays were performed using a monochromator microplate reader (Tecan M200 Pro). Complex I activity (NADH:ubiquinone oxidoreductase) was determined by measuring oxidation of NADH at 340 nm (using ferricyanide as the electron acceptor) in a reaction mixture of 50 mM potassium phosphate (pH 7.5), 0.2 mM NADH, and 1.7 mM potassium ferricyanide. Complex II activity (Succinate Dehydrogenase) was determined by measuring the reduction of the artificial electron acceptor 2,6-dichlorophenol-indophenol (DCIP) at 600 nm in a reaction mixture of 50 mM potassium phosphate (pH 7.5), 20 mM succinate, 2 μM DCIP, 10 μM rotenone, and 1 mM potassium cyanide. Complex III activity (Ubiquinol:cytochrome *c* oxidoreductase) was determined by measuring the reduction of cytochrome *c* at 550 nm in a reaction mixture of 50 mM potassium phosphate (pH 7.5), 35 μM reduced decylubiquinone, 15 μM cytochrome *c*, 10 μM rotenone, and 1 mM potassium cyanide. Complex IV activity (Cytochrome *c* oxidase) was determined by measuring the oxidation of cytochrome *c* at 550 nm in a reaction mixture of 50 mM potassium phosphate (pH 7.0) and 100 μM reduced cytochrome *c*. Citrate synthase activity was determined by measuring the reduction of 5,5’-dithiobis (2-nitrobenzoic acid) (DTNB) at 412 nm which was coupled to the reduction of acetyl-CoA by citrate synthase in the presence of oxaloacetate. The reaction mixture consisted of 100 mM Tris-HCl (pH 8.0), 100 μM DTNB, 50 μM acetyl-CoA, and 425 μM oxaloacetate. All activities were calculated as nmoles/min/mg protein, normalized to citrate synthase (CS) activity and finally expressed as the percentage of wild-type activity.

### Mitochondria Isolation

Freshly dissected BAT tissue from about 8-week-old male and female mice was homogenized in a Dounce homogenizer with 5ml ice-cold mitochondria isolation buffer (210mM Mannitol, 70mM Sucrose, 1mM EGTA, 5mM HEPES pH7.5, 0.5% BSA). The homogenates were filtered through cheesecloth to remove residual particulates and intact mitochondria were isolated by differential centrifugation using a previously described protocol ^63^. The mitochondrial pellet was resuspended in 25μL of isolation buffer and protein was quantitated using the Bradford assay (BioRad, #500-0006).

### Mass spectrometry

Purified BAT mitochondria from 10-12-week old male mice housed at RT or 30°C (n=3 for each genotype/condition) were resuspended in 8 M urea, 50 mM Tris, 5 mM CaCl_2_, 100 mM NaCl, and protease inhibitors. Mitochondria were lysed by probe sonication on ice, and proteins reduced by the addition of 5 mM DTT for 30 min at 37°C, followed cysteine alkylation by the addition of 15 mM iodoacetamide at RT for 45 min in the dark. The reaction was then quenched by the addition of 15 mM DTT for 15 minutes at RT. Proteins were first digested by the addition of endoproteinase LysC (Wako LC) at a 1:50 substrate:enzyme and incubated for 2h at RT. Next, samples were further digested by the addition of trypsin (Promega) at 1:100 substrate:enzyme, and incubated overnight at 37°C. Protein digests were then acidified by the addition of 0.5% triflororacetic acid, and samples desalted on C18 stage tips (Rainin). Peptides were resuspended in 4% formic acid and 3% acetonitrile, and approximately 1μg of digested mitochondria proteins was loaded onto a 75μm ID column packed with 25cm of Reprosil C18 1.9μm, 120Å particles. Peptides were eluted into an Orbitrap Fusion Tribrid (Thermo Fisher) mass spectrometer by gradient elution delivered by an Easy1200 nLC system (Thermo Fisher). The gradient was from 4.5% to 31% acetonitrile over 120 minutes. MS1 spectra were collected with oribitrap detection, while the 15 most abundant ions were fragmented by HCD and detected in the ion trap. All data were searched against the *Mus musculus* uniprot database (downloaded July 22, 2016). Peptide and protein identification searches were performed using the MaxQuant data analysis algorithm, and all peptide and protein identifications were filtered to a 1% false-discovery rate ^64, 65^. Label free quantification analysis was pefromed using the MSstats R-package ^66^. Proteome changes of each ETC complex were calculated by averaging log2 values of fold change of all identified proteins within individual ETC complex.

### Histology

Tissues were fixed in 10% formalin and processed and stained at AML Laboratories. Cell size was measured using ImageJ. Adipocyte size distribution was calculated using total adipocyte numbers counted in multiple images.

### Immunoblots

Puromycin (ThermoFisher, #A1113803) was injected intraperitoneally at the dose of 0.04 μmol g^-1^ 30 minutes prior to tissue collection. For lysates, tissues were lysed in ice-cold lysis-buffer (50 mM Tris-HCl, 150 mM NaCl, 1 mM EDTA, 6 mM EGTA, 20 mM NaF, 1% Triton X-100, 1μM MG132 and protease inhibitors) using a TissueLyser II (Qiagen). After centrifugation at 13000 rpm for 15 min, supernatants were reserved for protein determinations and SDS-PAGE analysis. Mitochondria were lyzed in the above lysis buffer before immunoblotting. Antibodies used were: Ucp1 (Sigma, #U6382), FLAG (Sigma, #F1804), Lrpprc (Santa Cruz Biotechnology, #SC-66844), Atf4 (Cell Signaling Technology, #11815), p-eIF2α (Cell Signaling Technology, #3398), eIF2α (Cell Signaling Technology, #5324), p-S6 (Cell Signaling Technology, #5364), S6 (Cell Signaling Technology, #2217X), p-4Ebp1 (Cell Signaling Technology, # 2855), 4Ebp1 (Cell Signaling Technology, #9452), Hsp90 (Santa Cruz Biotechnology, #SC-7949), total OXPHOS protein (Abcam, #ab110413), mt-Co2 (Proteintech, #55070-1-A), Cox4 (Cell Signaling Technology, #4850), Cox5b (Bethyl, #A-305-523A), Cox6b (Abgent, #AP20624a), Hsp60 (Bethyl, #A302-846A), puromycin (Kerafast, #EQ0001), and Ubiquitin (Santa Cruz Biotechnology, #SC-8017).

### Q-PCR and RNA-seq

Total RNA was extracted from tissues homogenized in TRIsure (Bioline, #BIO-38033) reagent and ISOLATE II RNA Mini kit (Bioline, #BIO-52073). Isolated RNA was reverse transcribed using iScript cDNA Synthesis Kit (Biorad, #170-8891), and the resulting cDNA was used for quantitative PCR on a CFX384 real-time PCR detection system (Bio-Rad). Relative mRNA expression level was determined using the 2(-Delta Ct) method with 36B4 as the internal reference control. Primer sequences are listed in Supplementary Table 2. RNA-seq was performed by Novogene Inc. Briefly, first strand cDNA was synthesized using random hexamer primer and M-MuLV Reverse Transcriptase (RNase H). Second strand cDNA synthesis was subsequently performed using DNA Polymerase I and RNase H. Double-stranded cDNA was purified using AMPure XP beads. Remaining overhangs of the purified double-stranded cDNA were converted into blunt ends via exonuclease/polymerase activities. After adenylation of 3’ ends of DNA fragments, NEBNext Adaptor with hairpin loop structure was ligated to prepare for hybridization. In order to select cDNA fragments of preferentially 150∼200bp in length, the library fragments were purified with AMPure XP system (Beckman Coulter, Beverly, USA). The libraries were sequenced in Illumina for 20 million reads with pair-end 150 bp (PE150). Downstream analysis was performed using a combination of programs including STAR, HTseq and Cufflink. Alignments were parsed using Tophat program and differentially expressed genes (DEGs) were determined through DESeq2/edgeR. KEGG enrichment was implemented by ClusterProfiler. Cis-regulatory sequence analysis was performed using iRegulon plugin in Cytoscape.

### mtDNA Quantification

The relative mtDNA content was measured using qPCR. The β2 microglobulin gene (B2M) was used as the nuclear gene (nDNA) normalizer for calculation of the mtDNA/nDNA ratio. A 322bp region of the mouse mtDNA was amplified using forward primer mtDNAF (CGACCTCGATGTTGGATCA) and the reverse primer mtDNAR (AGAGGATTTGAACCTCTGG). A fragment of the B2M gene was amplified using forward primer, B2MF (TCTCTGCTCCCCACCTCTAAGT), and reverse primer, B2MR (TGCTGTCTCGATGTTTGATGTATCT), giving an amplicon of 106 bp. The relative mtDNA content was calculated using the formula: mtDNA content = 1/2^ΔCt^, where 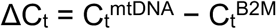.

### Quantification and statistical analysis

Data was presented as average ± SEM. Statistical significance was determined by t-test using GraphPad Prism 7. *: *p*<0.05 and **: *p*<0.01. Sample sizes for animal experiments were selected based on numbers typically used in similar published studies. No randomization of animals or predetermination of sample sizes by statistical methods was performed. No samples were measured repeatedly. *In vivo* metabolic experiments were repeated 2-3 times.

### Data accessibility

The mass spectrometry data files (raw and search results) have been deposited to the ProteomeXchange Consortium (http://proteomecentral.proteomexchange.org) via the PRIDE partner repository with dataset identifier PXD008798. The raw RNA-seq data has been deposited to NCBI GEO (accession number GSE117985).

## Acknowledgements

This work is supported by National Institutes of Health (NIH) grants DK105175 (B.W.), U54NS100717 (N.J.K.), P50GM082250 (N.J.K.), UCSF Diabetes Research Center P30DK063720 (B.W.), UCSF Nutrition Obesity Research Center P30DK098722 (B.W.), S10RR023051 (S.Y.). E.P. is supported by a fellowship grant from Hillblom foundation. We would like to thank Christophe Paillart and Vassily Kutyavin for assistance with indirect calorimetry experiments.

## Author contributions

B.W. and E.P. planned the experiments and wrote the manuscript. E.P. performed and analyzed thermogenic and metabolic phenotypes in animal studies. Y.Z. assisted in mouse colony maintenance, immunoblots and various assays. R.M. participated the initial studies. D.L.S., D.J-M., M.S., and N.J.K. performed mass spectrometry experiment and analyzed the data. T.L.H., and Y.S. performed ^18^F-FDG experiment.

## Competing interests

Authors declare no competing financial interests.

