## Supplementary data for "Brown adipocyte NOSEMPE promotes nonmitochondrial thermogenesis and improves systemic metabolism through ATF4 activation"

#### Supplementary Information

Supplementary material includes 15 supplementary figures and 2 supplementary tables.

**Supplementary Figure 1. (a)** Immunoblots of Lrpprc and Hsp90 in various tissues from ~8-week-old male CON and Lrpprc$^{BKO}$ mice. **(b)** Q-PCR analysis of mRNA levels of *Lrpprc* in BAT, iWAT and eWAT from ~8-week-old CON and Lrpprc$^{BKO}$ mice. **(c)** Immunoblots of Lrpprc, Ucp1 and Hsp90 in BAT from ~8-week-old CON and Lrpprc$^{BKO}$ mice. **(d)** Q-PCR analysis of relative mRNA levels of *Lrpprc, Ucp1,* $Pgc1\alpha$*, Cidea, Cox8b* and *Dio2* in BAT from ~8-week-old CON and Lrpprc$^{BKO}$ mice. Sample size: CON (n=4) and Lrpprc$^{BKO}$ (n=4). **(e)** Relative mtDNA copy numbers in BAT from ~8-week-old CON and Lrpprc$^{BKO}$ mice. Sample size: CON (n=5) and Lrpprc$^{BKO}$ (n=5). Data was presented as average ± SEM. Student t-test. *: p<0.05 and **: p<0.01.

**Supplementary Figure 2. (a)** Frequency of mitochondrial and non-mitochondrial proteins identified by mass spectrometry from isolated mitochondria of ~8-10-week old male CON and Lrpprc$^{BKO}$ mice housed at RT and 30°C. Sample size: n=3 for each condition (CON/RT, CON/30°C, Lrpprc$^{BKO}$/RT and Lrpprc$^{BKO}$/30°C). **(b)** Volcano plots showing significantly (p<0.1) down- or up-regulated mitochondrial proteins (over 1.5-fold) in Lrpprc$^{BKO}$ mice at RT and 30°C. **(c)** Principle component analysis of mitochondrial proteome in the four conditions. **(d)** Heatmap of Log2 FC values of the complex IV protein abundances measured by mass spectrometry in BAT mitochondria from CON and Lrpprc$^{BKO}$ mice housed at RT and 30°C. mt-unit: mitochondrion-encoded subunit, n-unit:

nucleus-encoded subunit, n-af: nucleus-encoded assembly factor. **(e)** Gene Ontology analysis showing enriched cellular components and molecular functions in mitochondrial proteome. **(f)** Heatmaps of log2FC of proteins important for Glycolysis, TCA cycle and beta-oxidation in Lrpprc$^{BKO}$ mice at RT and 30°C. **(g)** Mitochondrial cocktail immunoblot showing amounts of representative protein abundance of each ETC complex, Ndufb8 (complex I), Sdhb (complex II), Uqcrc2 (complex III), mt-Co1 (complex IV) and Atp5a (complex V), of BAT from ~8-12-week-old male CON and Lrpprc$^{BKO}$ mice housed at RT and 30°C. **(h)** Clustering analysis of mitochondrial proteome using fold changes of BAT mitochondrial protein abundance from ~8-12-week-old male Lkb1$^{BKO}$, Tfam$^{BKO}$ and Lrpprc$^{BKO}$ (to their relative controls) on normal chow at both RT and 30ºC. **(i)** Lists of proteins that are both upregulated or downregulated in the BAT from Tfam$^{BKO}$ and Lrpprc$^{BKO}$ mice.

**Supplementary Figure 3.** Recordings of energy expenditure (EE, calculated from oxygen consumption) in ~8-12-week old male CON and Lrpprc$^{BKO}$ mice for three days at RT **(a)** and 30°C **(b)**. Red arrowhead: time of CL injection. Recordings of respiratory exchange ratio (RER) in ~8-12-week old male CON and Lrpprc$^{BKO}$ mice for three days at RT **(c)** and 30°C **(d)**. Red arrowhead: time of CL injection. Average night and day RER **(e)**, food intake **(f)** and physical activity **(g)** from above mice. Sample size: CON/RT (n=6), Lrpprc$^{BKO}$/RT (n=6), CON/30°C (n=6) and Lrpprc$^{BKO}$/30°C (n=7). Data was presented as average ± SEM. Student t-test. *: $p<0.05$ and **: $p<0.01$.

**Supplementary Figure 4. (a)** Representative Sagittal views of fused PET and CT, and PET-only showing $^{18}$F-FDG uptake in BAT from CON and Lrpprc$^{BKO}$ mice. Color map showed at the right. **(b)** Average BAT $^{18}$F-FDG uptake in CON and Lrpprc$^{BKO}$ mice. Sample size: CON (n=5) and Lrpprc$^{BKO}$ (n=3). Data was presented as average ± SEM. Student t-test. *: p<0.05 and **: p<0.01.

**Supplementary Figure 5. (a)** Venn diagram showing down-regulated DEGs in the BAT of Lrpprc$^{BKO}$ mice at RT and 30°C. **(b)** GO analysis of shared down-regulated DEGs. **(c)** KEGG mmu00190 Oxidative phosphorylation pathway. Down-regulated genes were highlighted in blue. The mtDNA-encoded ETC subunits were labeled by a red asterisk. **(d)** Venn diagram showing up-regulated DEGs in the BAT of Lrpprc$^{BKO}$ mice at RT and thermoneutrality. **(e)** GO analysis of shared up-regulated DEGs.

**Supplementary Figure 6. (a)** Heatmap showing log2 fold changes of known ATF4 target genes in the BAT of Lrpprc$^{BKO}$ mice at normal chow or after 4-week HFD at both RT and 30ºC. Sample size: CON/RT/NC (n=8), Lrpprc$^{BKO}$/RT/NC (n=8), CON/30°C/NC (n=4), Lrpprc$^{BKO}$/30°C/NC (n=6), CON/RT/HFD (n=8), Lrpprc$^{BKO}$/RT/HFD (n=7), CON/30°C/HFD (n=4) and Lrpprc$^{BKO}$/30°C/HFD (n=6). Immunoblots showing amounts of Lrpprc, p-eIF2$\alpha$, total eIF2$\alpha$, and Hsp90 in the BAT of ~8-12-week-old male Tfam$^{BKO}$ **(b)** and betaless **(c)** mice compared with their relative controls at normal chow at both RT and 30ºC.

**Supplementary Figure 7. (a)** Cross scheme to generate brown adipocyte-specific ATF4 overexpression **(**ATF4$^{BOX}$) and its control (CON). **(b)** Flag immunoblots of Flat-ATF4 and Hsp90 in the BAT from ~8-week-old male CON and ATF4$^{BOX}$ mice at RT. **(b)** Q-PCR analysis of mRNA levels of thermogenic genes (*Ucp1, Cidea, Cox8b and Pgc1$\alpha$)* in the BAT of ~8-week-old CON and ATF4$^{BOX}$ mice at RT. Sample size: CON (n=5) and ATF4$^{BOX}$ (n=5). Average night and day RER **(d)**, food intake **(e)** and physical activity **(f)** in ~8-12-week old male CON and ATF4$^{BOX}$ mice for three days at RT and 30°C. Sample size: CON/RT (n=10), ATF4$^{BOX}$/RT (n=4), CON/30°C (n=10) and ATF4$^{BOX}$/30°C (n=5). Data was presented as average ± SEM.

**Supplementary Figure 8.** Immunoblots of p-S6, total S6, p-4Ebp1, total 4Ebp1, puromycylated protein, ubiquitinated protein and Hsp90 in the BAT **(a)** and muscle **(b)** of ~10-week-old male CON and ATF4$^{BOX}$ mice housed at 30ºC with DMSO or rapamycin treatment.

**Supplementary Figure 9.** Immunoblots of p-S6, total S6, p-4Ebp1, total 4Ebp1, puromycylated protein, ubiquitinated protein and Hsp90 in the BAT of ~10-week-old male Lrpprc$^{BKO}$ mice **(a)** and Tfam$^{BKO}$ mice **(b)** at RT and 30ºC

**Supplementary Figure 10. (a)** Body weight of male CON and Lrpprc$^{BKO}$ mice at RT and 30°C fed with normal chow (NC). **(b)** Body weight, lean mass, fat mass, and fat percentage of ~8-month-old male CON and Lrpprc$^{BKO}$ mice at RT and 30°C. **(c)** Tissue mass of BAT, iWAT, and eWAT of ~8-month-old male CON and Lrpprc$^{BKO}$ mice. Sample size: CON/RT (n=8), Lrpprc$^{BKO}$/RT (n=10), CON/30ºC (n=7) and Lrpprc$^{BKO}$/30ºC (n=5). Data was presented as average ± SEM. Student t-test. *: p<0.05 and **: p<0.01.

**Supplementary Figure 11. (a)** Body weight of female CON and Lrpprc$^{BKO}$ mice after 12-week HFD at RT and 30°C. Sample size: CON/RT (n=8), Lrpprc$^{BKO}$/RT (n=7), CON/30°C (n=11) and Lrpprc$^{BKO}$/30°C (n=17). **(b)** Body weight, lean mass, fat mass, and fat percentage of female CON and Lrpprc$^{BKO}$ mice after 12-week HFD. Sample size: CON/RT (n=8), Lrpprc$^{BKO}$/RT (n=7), CON/30°C (n=8) and Lrpprc$^{BKO}$/30°C (n=11). **(c)** Tissue mass of eWAT, iWAT, and BAT of female CON and Lrpprc$^{BKO}$ mice before and after HFD**.** Sample size: CON/RT (n=8), Lrpprc$^{BKO}$/RT (n=7), CON/30°C (n=18) and Lrpprc$^{BKO}$/30°C (n=17). **(d)** Liver triglyceride contents of female CON and Lrpprc$^{BKO}$ mice after 12-week HFD. Sample size: CON/RT (n=8), Lrpprc$^{BKO}$/RT (n=7), CON/30°C (n=8) and Lrpprc$^{BKO}$/30°C (n=11). Data was presented as average ± SEM. Student t-test. *: p<0.05 and **: p<0.01.

**Supplementary Figure 12. (a)** Representative H&E staining of BAT from ~8-12-week of male CON and Atf4$^{BKO}$ mice housed at RT and 30°C. Scale bar: 50μm. **(b)** Heatmap showing log2 fold changes of mtDNA- and nuclear-encoded ETC genes, known ATF4 target genes in the BAT of CON and Atf4$^{BKO}$ mice housed at RT and 30°C. Hourly CL-induced EE **(d),** average night and day EE **(c)**, RER **(e)**, food intake **(f)** and physical activity **(g)** in ~8-12-week old male CON and ATF4$^{BOX}$ mice for three days at RT and 30°C. Sample size: CON/RT (n=4), Atf4$^{BKO}$/RT (n=3), CON/30°C (n=6) and Atf4$^{BKO}$/30°C (n=6). **(h)** Body weight of male CON and Atf4$^{BKO}$ mice under 12-week HFD. Sample size:

CON/RT (n=10), Atf4$^{BKO}$/RT (n=10), CON/30°C (n=13) and Atf4$^{BKO}$/30°C (n=13). **(i)** Body weight, lean mass, fat mass, and fat percentage of male CON and Atf4$^{BKO}$ mice under 12-week HFD. **(j)** Tissue mass of BAT, iWAT, and eWAT of male CON and Atf4$^{BKO}$ mice under 12-week HFD. **(k)** Serum glucose levels during ITT in male CON and Atf4$^{BKO}$ mice after 12-week HFD. **(l)** Area under the curve (AUC) values of glucose levels in ITTs showed. Sample size: CON/RT (n=5), Atf4$^{BKO}$/RT (n=8), CON/30°C (n=16) and Atf4$^{BKO}$/30°C (n=8). Serum insulin **(m),** serum triglyceride contents **(n)** and liver triglyceride contents **(o)** of male CON, Lrpprc$^{BKO}$ and Lrpprc;Atf4$^{BKO}$ mice after HFD**.** Sample size: CON/RT (n=5), Atf4$^{BKO}$/RT (n=8), CON/30°C (n=8) and Atf4$^{BKO}$/30°C (n=8). Data was presented as average ± SEM.

**Supplementary Figure 13. (a)** Representative H&E staining of BAT from male CON, Lrpprc$^{BKO}$, Atf4$^{BKO}$, and Lrpprc;Atf4$^{BKO}$ mice housed at RT. Scale bar: 100 μm. **(b)** Heatmap showing log2 fold changes of mtDNA- and nuclear-encoded ETC genes, known ATF4 target genes in the BAT of Lrpprc$^{BKO}$ and Lrpprc;Atf4$^{BKO}$ mice housed at RT and 30°C. **(c)** CTT of ~8-12-week old male and female CON, Lrpprc$^{BKO}$, Atf4$^{BKO}$, and Lrpprc;Atf4$^{BKO}$ mice housed at RT. Sample size: CON (n=6), Lrpprc$^{BKO}$ (n=9), Atf4$^{BKO}$ (n=4), and Lrpprc;Atf4$^{BKO}$ (n=8). Data was presented as average ± SEM. Student t-test. **: $p<0.01$.

**Supplementary Figure 14.** Average night and day RER **(a)**, food intake **(b)** and physical activity **(c)** in ~10-week old male CON, Lrpprc$^{BKO}$ and Lrpprc;Atf4$^{BKO}$ mice at 30°C.

Sample size: CON (n=6), $Lrpprc^{BKO}$ (n=3), and $Lrpprc;Atf4^{BKO}$ (n=3). Data was presented as average ± SEM.

**Supplementary Figure 15. (a)** Body weight, lean mass, fat mass, and fat percentage of male CON, $Lrpprc^{BKO}$ and $Lrpprc;Atf4^{BKO}$ mice after 12-week HFD at 30°C. Sample size: CON (n=9), $Lrpprc^{BKO}$ (n=6), and $Lrpprc;Atf4^{BKO}$ (n=5). **(b)** Representative images of dissected iWAT, eWAT and BAT from male CON, $Lrpprc^{BKO}$ and $Lrpprc;Atf4^{BKO}$ mice after 12-week HFD. **(c)** Adipocyte size distribution in eWAT from male CON, $Lrpprc^{BKO}$ and $Lrpprc;Atf4^{BKO}$ mice after 12-week HFD. Total adipocytes counted: CON (n=681), $Lrpprc^{BKO}$ (n=983), and $Lrpprc;Atf4^{BKO}$ (n=945). **(d)** Representative H&E staining of eWAT and liver from male CON, $Lrpprc^{BKO}$ and $Lrpprc;Atf4^{BKO}$ mice after 12-week HFD. Scale bar: 50 μm. **(k)** Serum glucose levels during ITT in male CON, $Lrpprc^{BKO}$ and $Lrpprc;Atf4^{BKO}$ mice after 12-week HFD. **(l)** Area under the curve (AUC) values of glucose levels in ITTs showed. Sample size: CON (n=9), $Lrpprc^{BKO}$ (n=6), and $Lrpprc;Atf4^{BKO}$ (n=5). Data was presented as average ± SEM. Student t-test. *: $p<0.05$ and **: $p<0.01$.

**Supplementary Table 1.**

Excel table of mass spectrometry data of mitochondrial proteome from CON and $Lrpprc^{BKO}$ mice at RT and 30ºC.

https://ucsf.box.com/s/0b68suk8f7jtdj8u94c0w5gj4bcjvkf3

**Supplementary Table 2.**

List of q-PCR primers.

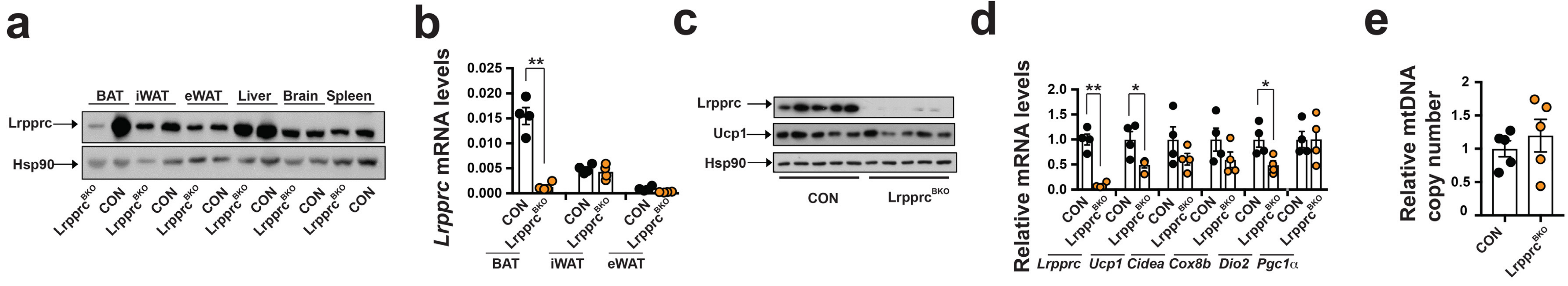


### Supplementary Figure 1

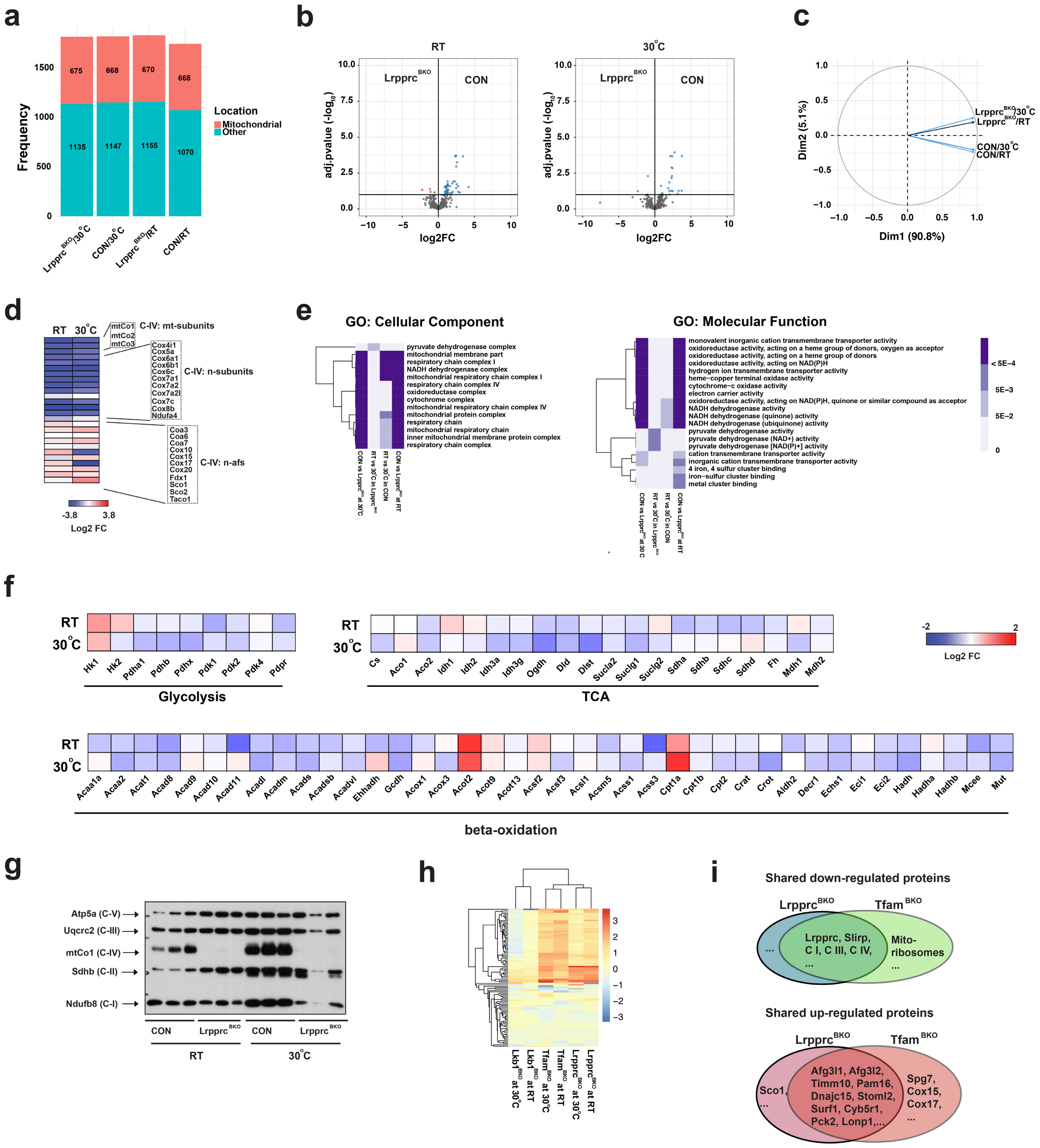


Supplementary Figure 2

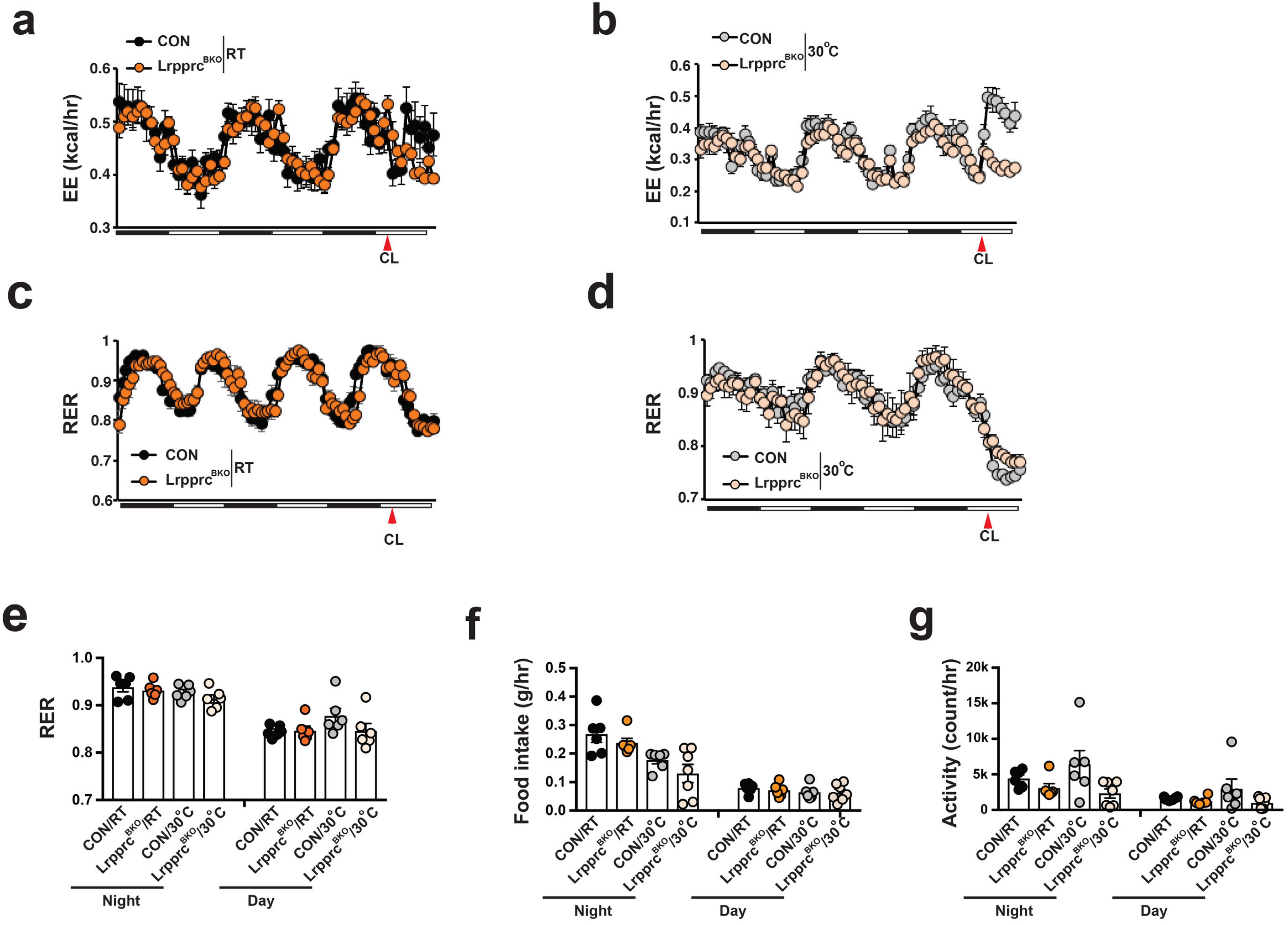


### Supplementary Figure 3

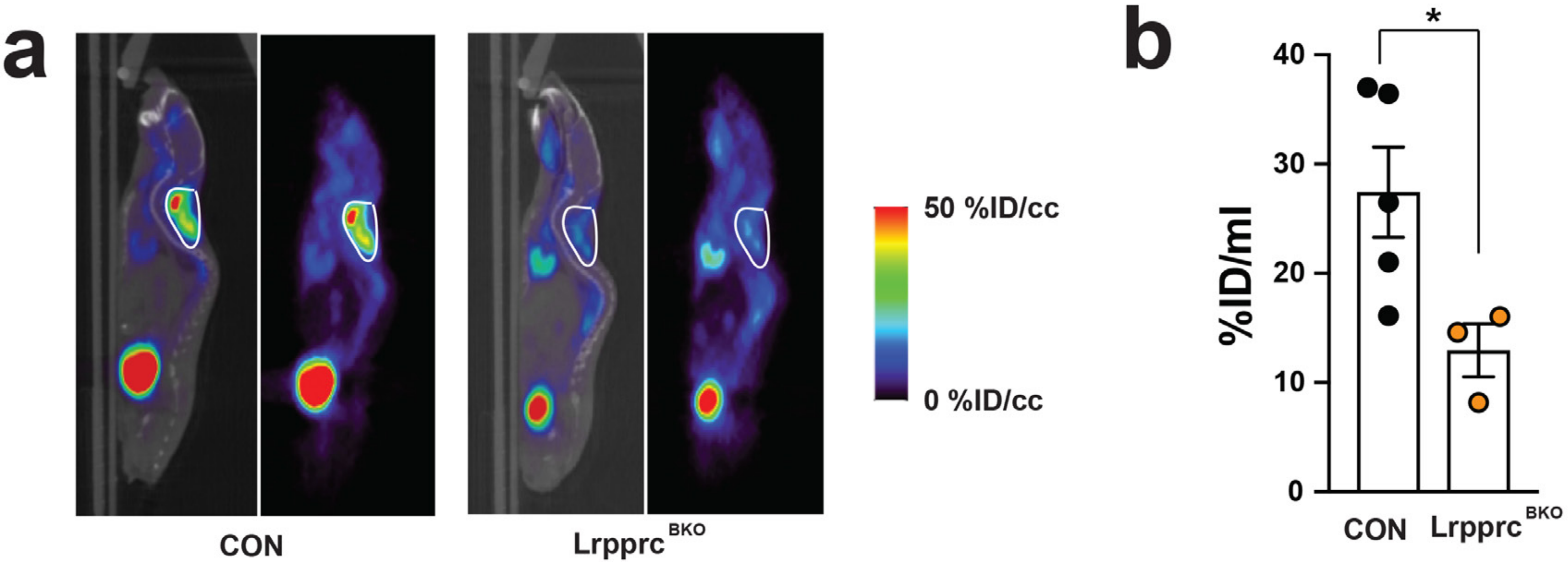


### Supplementary Figure 4

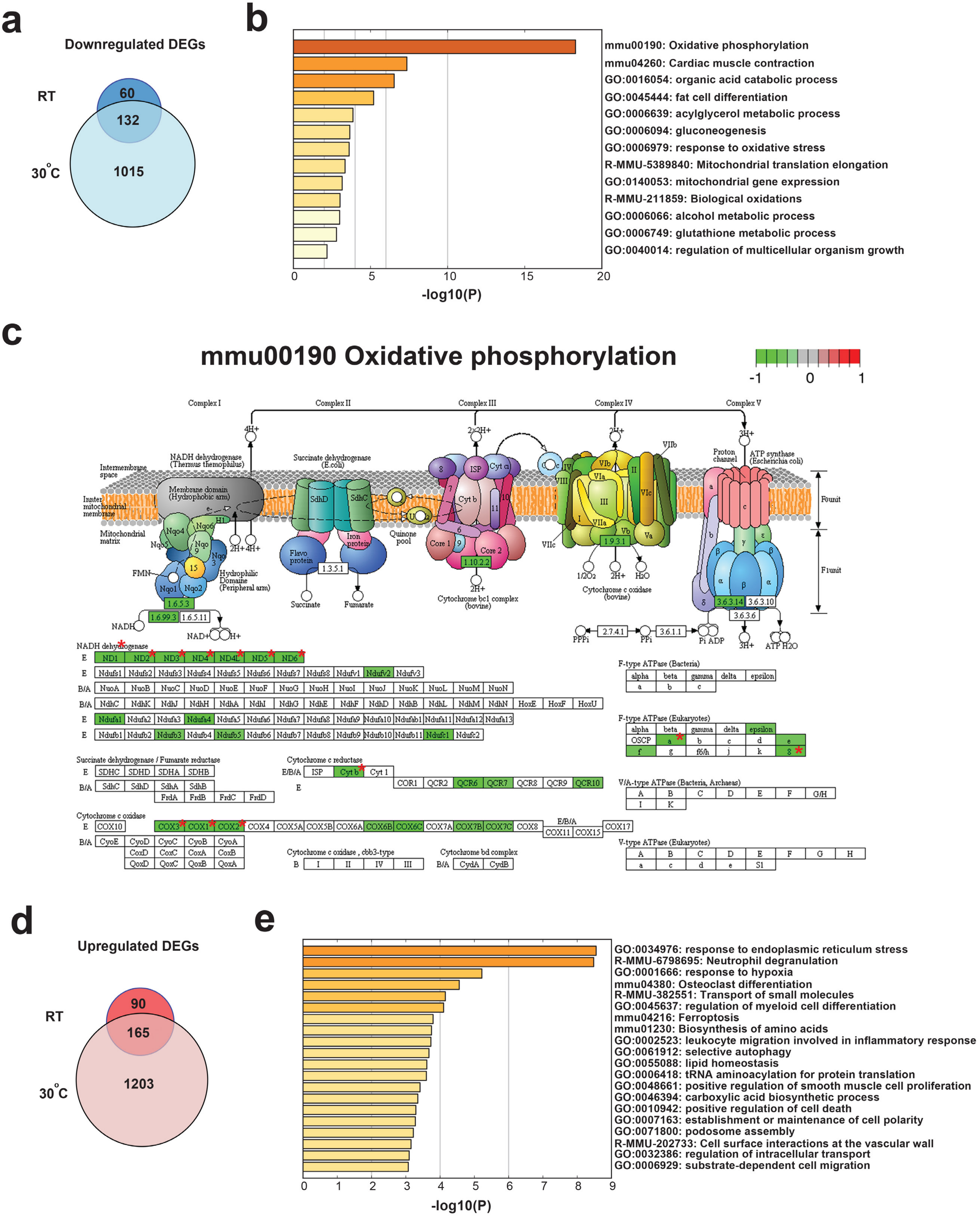


### Supplementary Figure 5

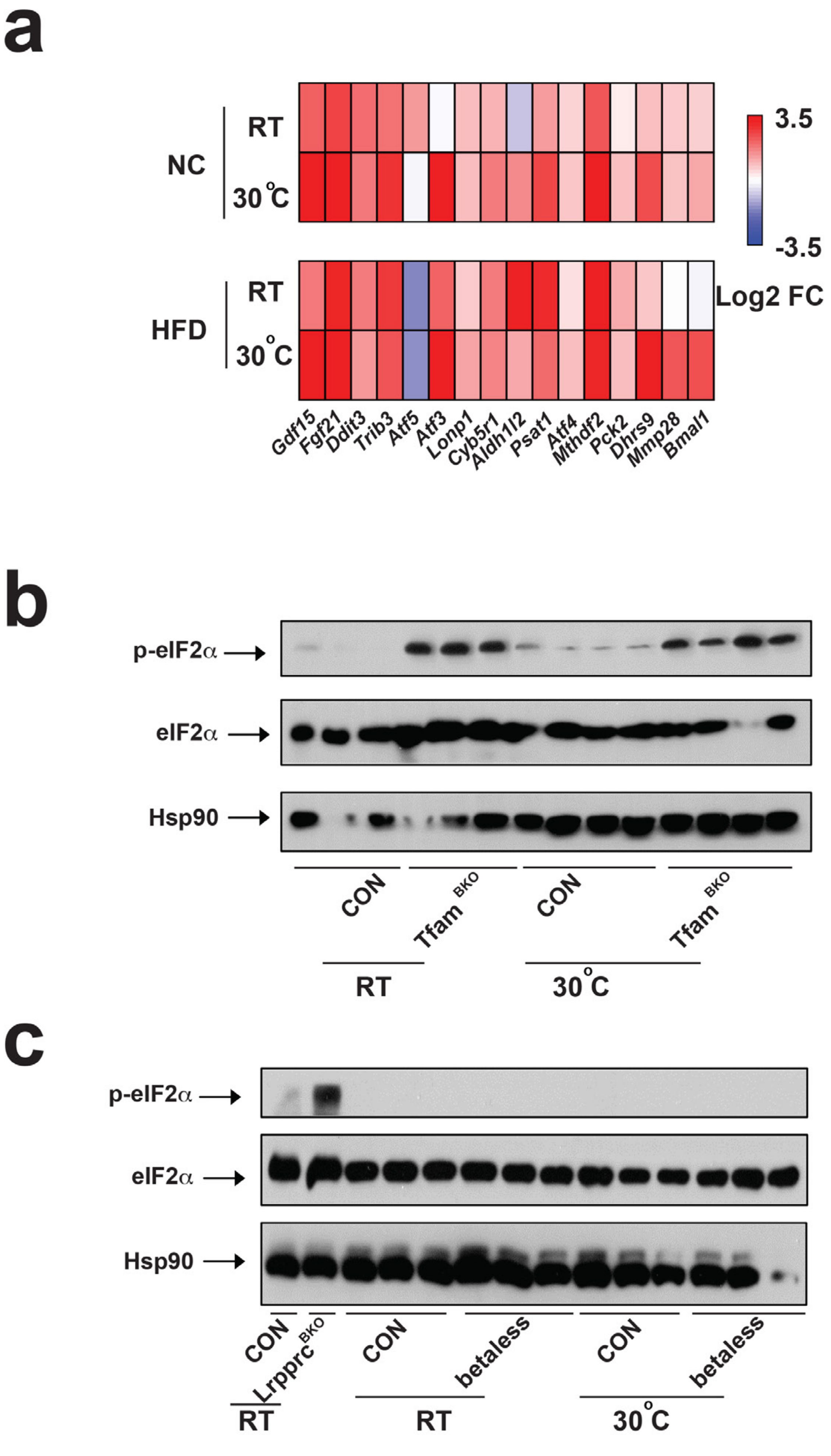


### Supplementary Figure 6

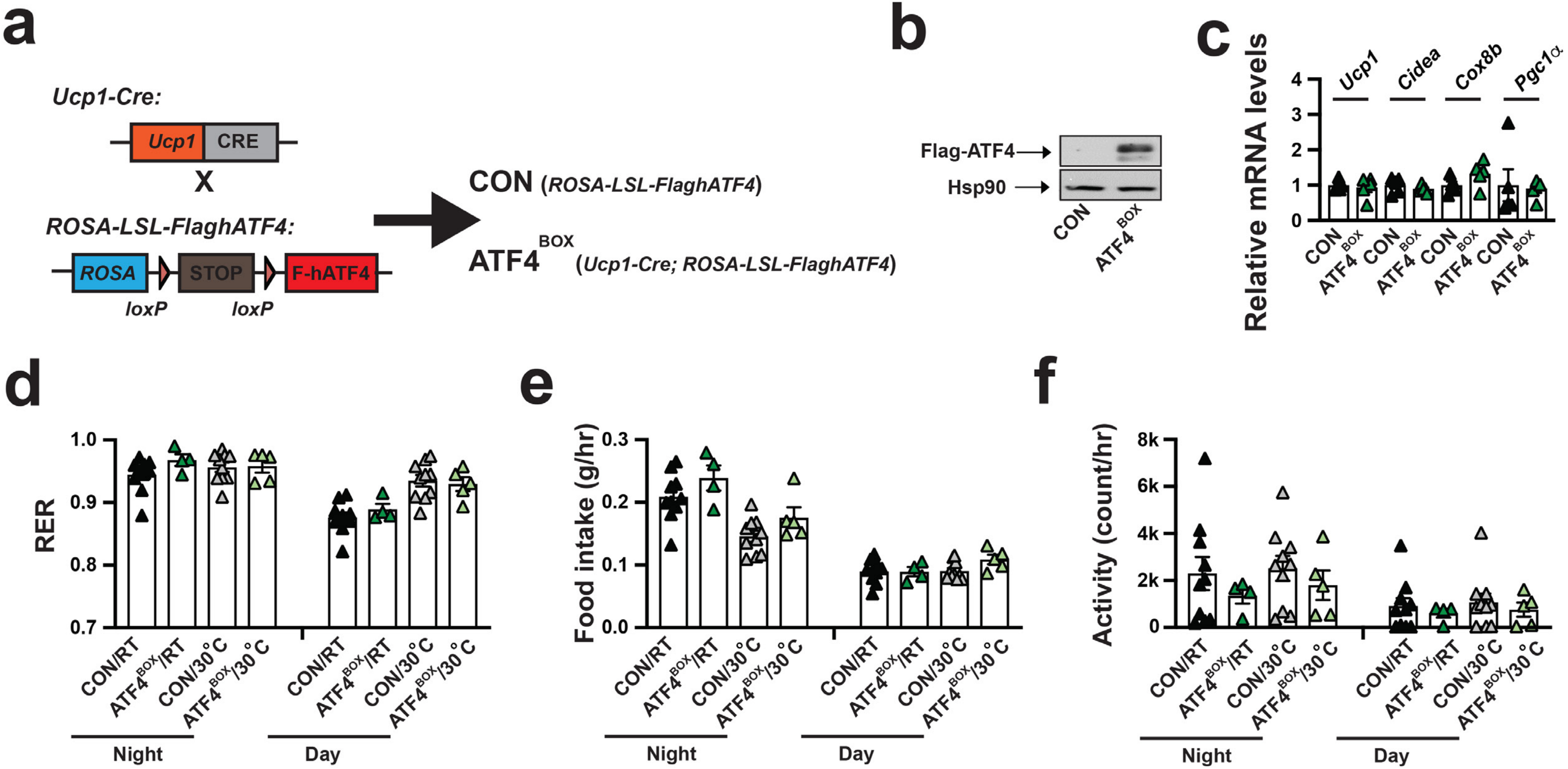


### Supplementary Figure 7

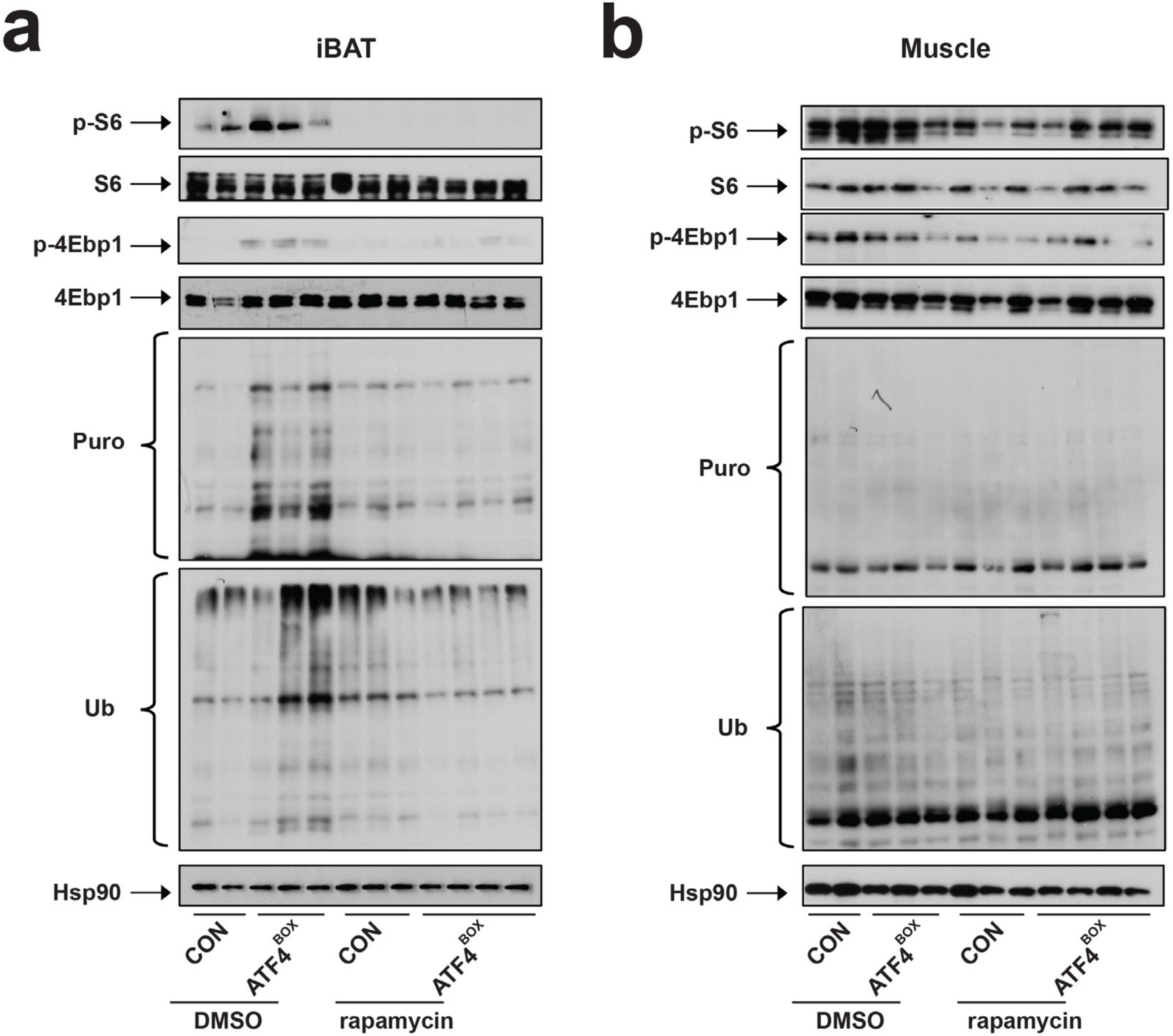


### Supplementary Figure 8

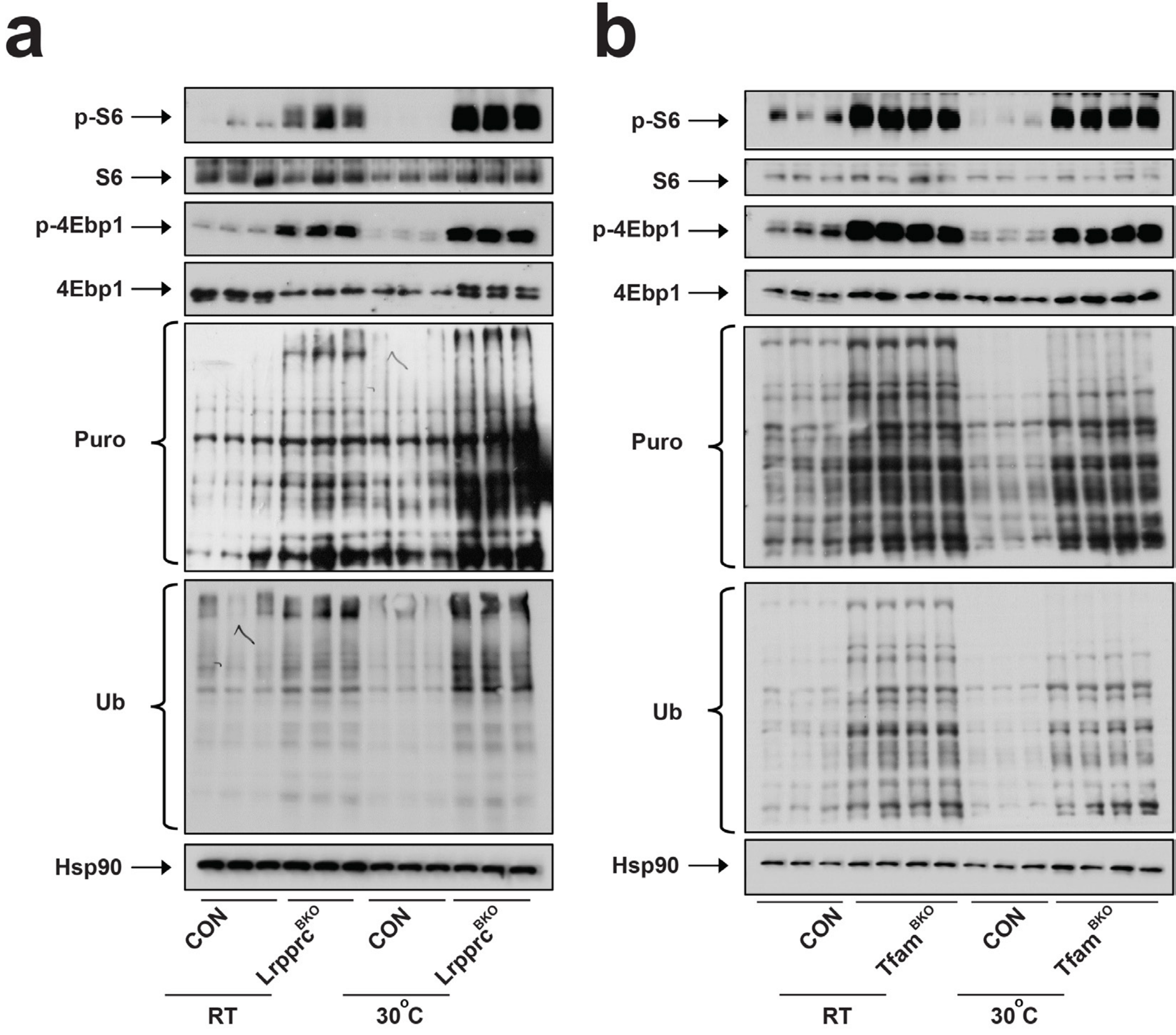


### Supplementary Figure 9

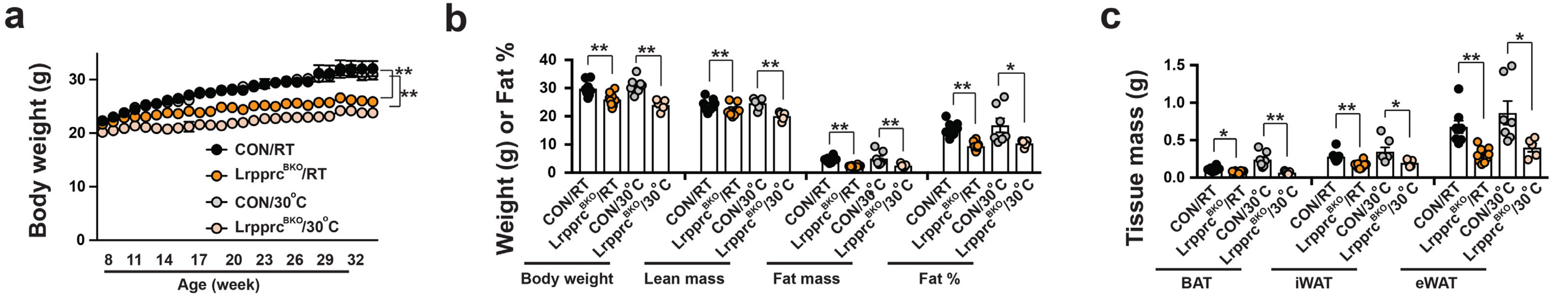


### Supplementary Figure 10

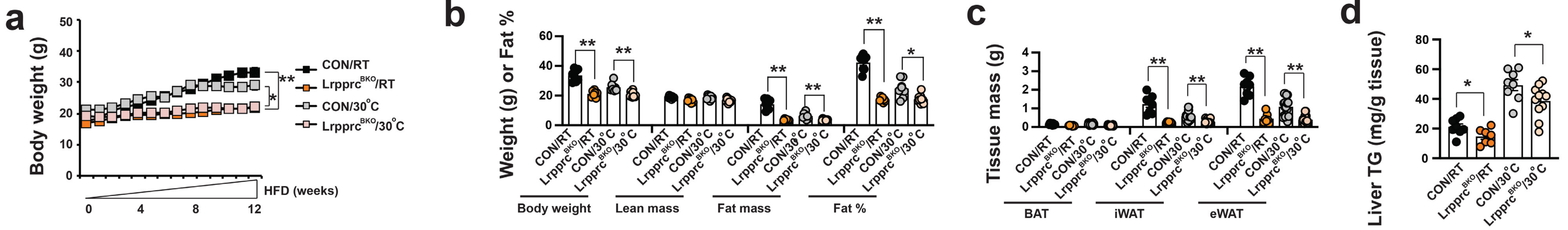


### Supplementary Figure 11

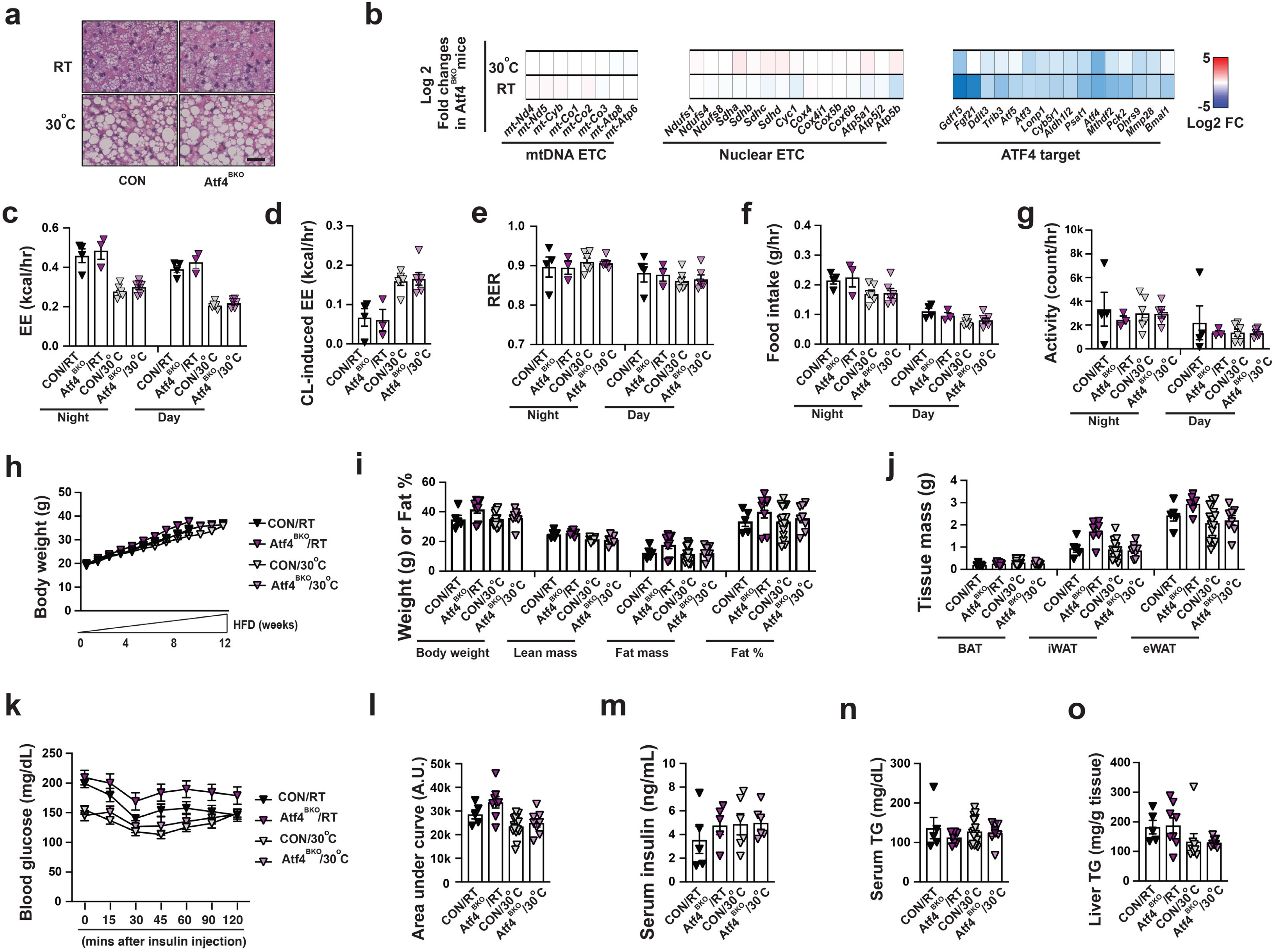


### Supplementary Figure 12

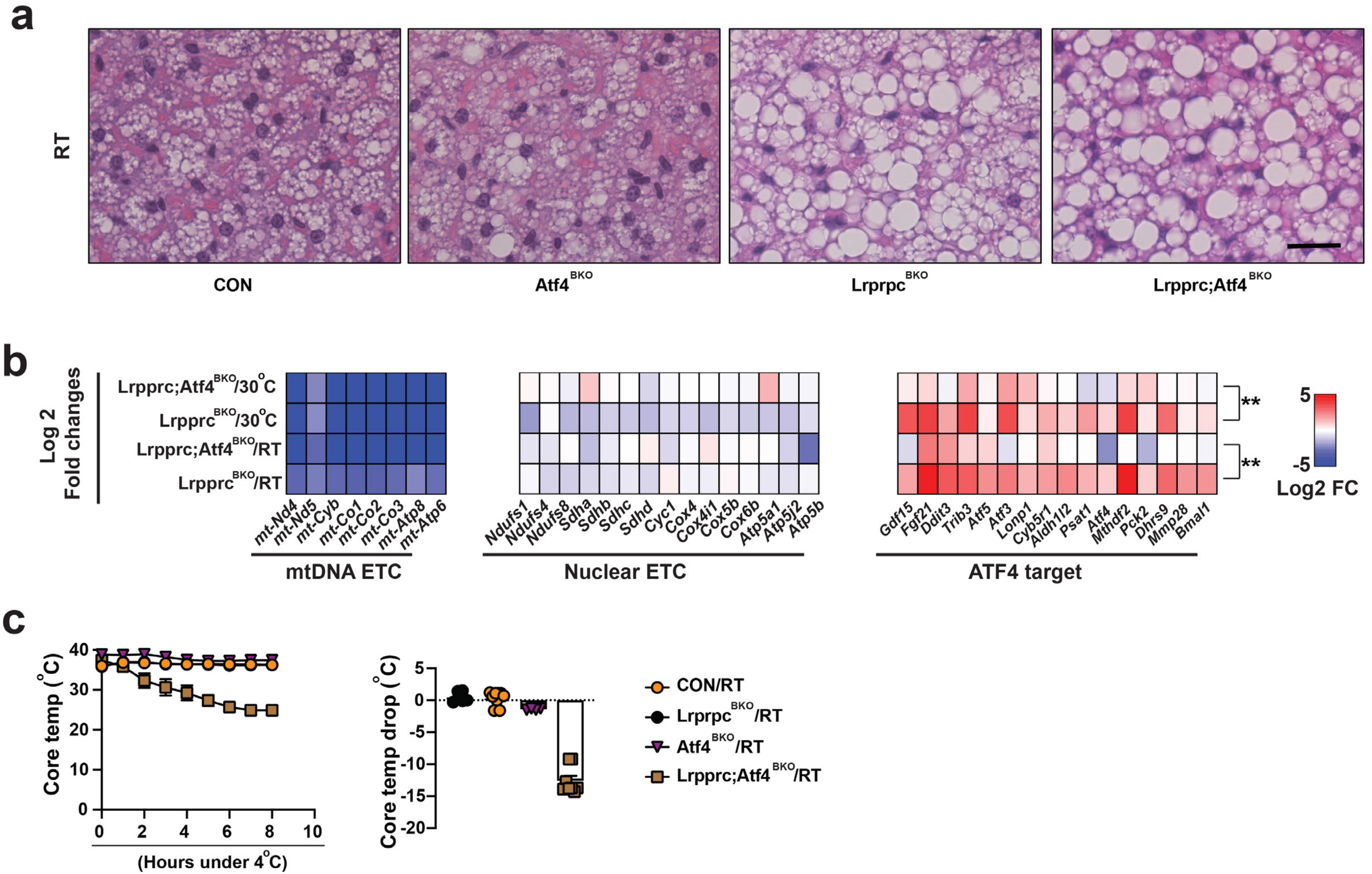


### Supplementary Figure 13

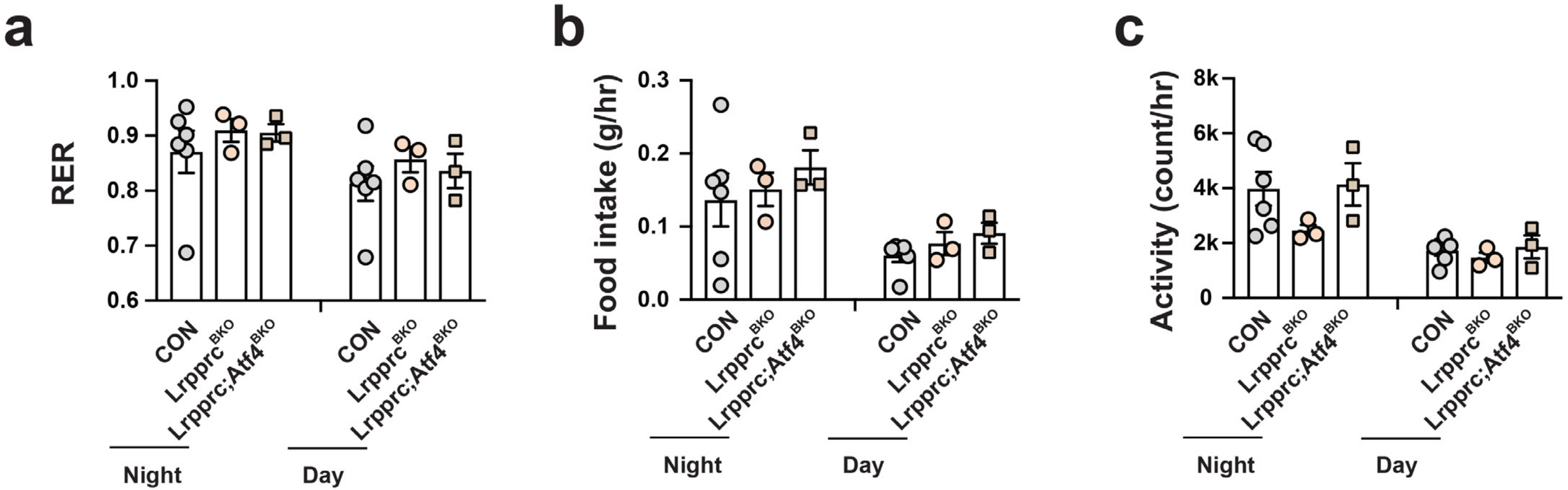


### Supplementary Figure 14

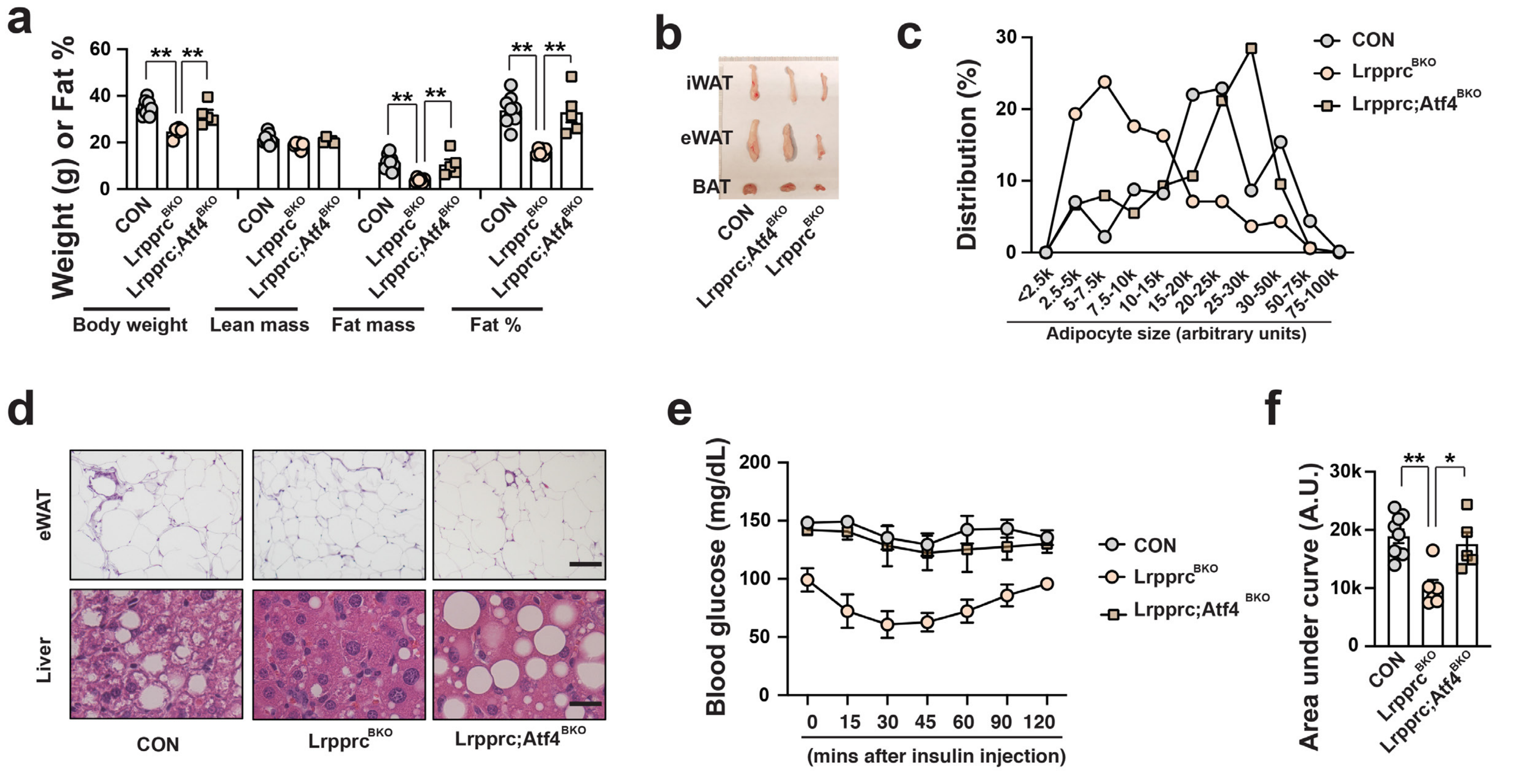


### Supplementary Figure 15

**Supplementary Table 2: List of primer sequences for q-PCR**

| | |
|---|---|
| 36B4-F | TTTGGGCATCACCACGAAAA |
| 36B4-R | GGACACCCTCCAGAAAGCGA |
| Lrpprc-F | AGCCTGCTCCTGTGAGAAAG |
| Lrpprc-R | TCCCAGATCTTGTGAGCAAA |
| mt-Nd4-F | CTAATAATCGCACATGGCCTC |
| mt-Nd4-R | CGTAGTTGGAGTTTGCTAGG |
| mt-Nd5-F | CATCCTTCTCAACTTTACTGGG |
| mt-Nd5-R | TTTATGGGTGTAATGCGGT |
| mt-Cyb-F | CCATTCTACGCTCAATCCCCA |
| mt-Cyb-R | AGGCTTCGTTGCTTTGAGGTA |
| mt-Co1-F | ACACAACTTTCTTTGATCCCG |
| mt-Co1-R | AGAATCAGAACAGATGCTGG |
| mt-Co2-F | ATAATCCCAACAAACGACCT |
| mt-Co2-R | CTCGGTTATCAACTTCTAGCA |
| mt-Co3-F | GGTATAATTCTATTCATCGTCTCGG |
| mt-Co3-R | AGAACGCTCAGAAGAATCCT |
| mt-Atp8-F | GGCACCTTCACCAAAATCACT |
| mt-Atp8-R | GGGGTAATGAATGAGGCAAATAGA |
| mt-Atp6-F | CCTTCAATCCTATTCCCATCC |
| mt-Atp6-R | GTTGGAAAGAATGGAGACGG |
| Ndufs1-F | AGGATATGTTCGCACAACTGG |
| Ndufs1-R | TCATGGTAACAGAATCGAGGGA |
| Ndufs4-F | CTGCCGTTTCCGTCTGTAGAG |
| Ndufs4-R | TGTTATTGCGAGCAGGAACAAA |
| Ndufs8-F | AGTGGCGGCAACGTACAAG |
| Ndufs8-R | TCGAAAGAGGTAACTTAGGGTCA |
| Sdha-F | GGAACACTCCAAAAACAGACCT |
| Sdha-R | CCACCACTGGGTATTGAGTAGAA |
| Sdhb-F | AATTTGCCATTTACCGATGGGA |
| Sdhb-R | AGCATCCAACACCATAGGTCC |
| Sdhc-F | GCTGCGTTCTTGCTGAGACA |
| Sdhc-R | ATCTCCTCCTTAGCTGTGGTT |
| Sdhd-F | TGGTCAGACCCGCTTATGTG |
| Sdhd-R | GGTCCAGTGGAGAGATGCAG |
| Cyc1-F | CAGCTTCCATTGCGGACAC |
| Cyc1-R | GGCACTCACGGCAGAATGAA |
| Cox4-F | ATGTCACGATGCTGTCTGCC |
| Cox4-R | GTGCCCCTGTTCATCTCGGC |
| Cox4i1-F | ATTGGCAAGAGAGCCATTTCTAC |
| Cox4i1-R | CACGCCGATCAGCGTAAGT |
| Cox5a-F | GGGTCACACGAGACAGATGA |
| Cox5a-R | CCAAGATGCGAACAGCACTA |
| Cox5b-F | GATGAGGAGCAGGCTACTGG |
| Cox5b-R | TGCAGCCCACTATTCTCTTG |

| | |
|---|---|
| Cox6b1-F | CCCCAACCAGAACCAGACTA |
| Cox6b1-R | GATCTTCCCAGGAAATGTGC |
| Atp5a1-F | TCTCCATGCCTCTAACACTCG |
| Atp5a1-R | CCAGGTCAACAGACGTGTCAG |
| Atp5j2-F | TGCCGAGCTGGATAATGATGC |
| Atp5j2-R | ACCATGCTAATCCCCGAGATG |
| Atp5b-F | GCAAGGCAGGGACAGCAGA |
| Atp5b-R | CCCAAGGTCTCAGGACCAACA |
| Gdf15-F | CTGGCAATGCCTGAACAACG |
| Gdf15-R | GGTCGGGACTTGGTTCTGAG |
| Fgf21-F | GTGTCAAAGCCTCTAGGTTTCTT |
| Fgf21-R | GGTACACATTGTAACCGTCCTC |
| Ddit3-F | CTGCCTTTCACCTTGGAGAC |
| Ddit3-R | GGACGCAGGGTCAAGAGTAG |
| Trib3-F | GGGGCCTTATATCCTTTTGG |
| Trib3-R | GCAGGGTACACCTTGCAG |
| Atf5-F | CCTTGCCCTTGCCACCTTTGAC |
| Atf5-R | CCAGAGGAGGAGGCTGCTGT |
| Atf3-F | GCTGCCAAGTGTCGAAACAAG |
| Atf3-R | CAGTTTTCCAATGGCTTCAGG |
| Lonp1-F | TGAGCTGCAAGATGTTCTGG |
| Lonp1-R | AGCCCCAATTCCTTCTTGAT |
| Cyb5r1-F | CTACCTCTCTGCCCGAATTG |
| Cyb5r1-R | CCCAATCTTCAGGCTATCCA |
| Aldh1l2-F | CACCCCTGTGATTGAGGACT |
| Aldh1l2-R | GCCTCTTCGTCCTCTCCTCT |
| Psat1-F | TGCTCGAAATGACTCACAGG |
| Psat1-R | CAGCACTCCTTCCAGCTTTC |
| Atf4-F | AAGGAGGAAGACACTCCCTCT |
| Atf4-R | CAGGTGGGTCATAAGGTTTGG |
| Mthdf2-F | AGGTCCCAAGCCTTTGAGTT |
| Mthfd2-R | GTAAGGGAGTGCCGTTGAAA |
| Pck2-F | ATGGCTGCTATGTACCTCCC |
| Pck2-R | GCGCCACAAAGTCTCGAAC |
| Dhrs9-F | TACCTCCTCGGTGAACTTGG |
| Dhrs9-R | TGGGATTTGCCAGCTCTACT |
| Mmp28-F | GAGGCGTAAGAAACGCTTTG |
| Mmp28-R | CCAGAACTCCAGTGCTGACA |
| Bmal1-F | TGACCCTCATGGAAGGTTAGAA |
| Bmal1-R | GGACATTGCATTGCATGTTGG |
| Ucp1-F | ACTGCCACACCTCCAGTCATT |
| Ucp1-R | CTTTGCCTCACTCAGGATTGG |
| Cox8b-F | GAACCATGAAGCCAACGACT |
| Cox8b-R | GCGAAGTTCACAGTGGTTCC |
| Cidea-F | TGCTCTTCTGTATCGCCCAGT |
| Cidea-R | GCCGTGTTAAGGAATCTGCTG |

| | |
|---|---|
| Dio2-F | CAGTGTGGTGCACGTCTCCAATC |
| Dio2-R | TGAACCAAAGTTGACCACCAG |
| Pgc1a-F | AGCCGTGACCACTGACAACGAG |
| Pgc1a-R | GCTGCATGGTTCTGAGTGCTAAG |
| Cd68-F | GCAGCACAGTGGACATTCAT |
| Cd68-R | TTGCATTTCCACAGCAGAAG |
| F4/80-F | TTTCCTCGCCTGCTTCTTC |
| F4/80-R | CCCCGTCTCTGTATTCAACC |
| Cd11c-F | CAGAACTTCCCAACTGCACA |
| Cd11c-R | TCTCTGAAGCTGGCTCATCA |
| Leptin-F | GAGACCCCTGTGTCGGTTC |
| Leptin-R | CTGCGTGTGTGAAATGTCATTG |
| Ccl2-F | CTTCTGGGCCTGCTGTTCA |
| Ccl2-R | CCAGCCTACTCATTGGGATCA |
